# The AUX1-AFB1-CNGC14 module establishes longitudinal root surface pH profile

**DOI:** 10.1101/2022.11.23.517700

**Authors:** Nelson BC Serre, Daša Wernerová, Pruthvi Vittal, Shiv Mani Dubey, Eva Medvecká, Adriana Jelínková, Jan Petrášek, Guido Grossmann, Matyáš Fendrych

## Abstract

Plant roots navigate in the soil environments following the gravity vector. Cell divisions in the meristem and rapid cell growth in the elongation zone propel the root tips through the soil. Actively elongating cells acidify their apoplast to enable cell wall extension by the activity of plasma membrane AHA H^+^-ATPases. The phytohormone auxin, central regulator of gravitropic response and root development, inhibits root cell growth, likely by rising the pH of the apoplast. However, the role of auxin in the regulation of the apoplastic pH gradient along the root tip is unclear. Here we show, by using an improved method for visualization and quantification of root surface pH, that the *Arabidopsis thaliana* root surface pH shows distinct acidic and alkaline zones, which are not primarily determined by the activity of AHA H^+^-ATPases. Instead, the distinct domain of alkaline pH in the root transition zone is controlled by a rapid auxin response module, consisting of the AUX1 auxin influx carrier, the AFB1 auxin co-receptor and the CNCG14 calcium channel. We demonstrate that the rapid auxin response pathway is required for an efficient navigation of the root tip.

## Introduction

Plants colonize soil by the growth of their roots. Because plant cells are non-motile, mechanically coupled by their cell walls and symplastically connected by plasmodesmata, the extent and direction of root growth is driven by the elongation of cells that can differ across the organ (Braidwood *et al*. 2014). The balance between cell proliferation and differentiation in the root apex is controlled by several phytohormone and peptide signaling pathways, among which auxin plays a specific role thanks to its long-distance intercellular transport that serves as the information carrier between distant plant tissues (Grieneisen *et al*. 2007; Wisniewska *et al*. 2006).

The root apex is divided in several zones fulfilling various roles in root development. At the tip of the root, the root cap covers and protects the meristem, and is the center of gravity perception. In the meristem, stem cells produce transit-amplifying cells that populate the root tip (Dolan *et al*. 1993; Campilho *et al*. 2006). In the model plant *Arabidopsis thaliana*, the root cap reaches up to the transition zone, where cells stop dividing and prepare for elongation; the transition zone is characterized by a distinct physiology and shows a high responsiveness to external factors (Verbelen *et al*. 2006). Further in the shootward direction, cells rapidly elongate in the elongation zone, propelling the root tip through the soil (Beemster and Baskin 1998). After reaching their final length, cells differentiate in the maturation zone, where the root hairs emerge to increase the absorptive root surface.

To elongate, cells must soften their cell walls, increase turgor pressure or both (Lintilhac 2014). Proton extrusion into the apoplast by the plasma membrane (PM) H^+^ ATPases is crucial for both cell wall softening and turgor maintenance. Acidic pH leads to activation of cell wall remodeling enzymes, and the H^+^ gradient at the PM drives the transport of most nutrients and ions, and as such is required for turgor pressure generation (Falhof *et al*. 2015). It is therefore not surprising that root zones of numerous species show a distinct pattern of surface proton fluxes and cell wall pH (Siao *et al*. 2020). In general, the tip of the root and the late elongation and maturation zones show outward H^+^ fluxes, while the transition zone and early elongation zones show inward H^+^ fluxes (Weisenseel and Meyer 1997). Several studies showed that this battery-like system, with positive charges flowing out of the maturation zone and entering at the transition zone, is mainly driven by H^+^ fluxes without completely excluding calcium and chloride ions (Weisenseel *et al*. 1979; Björkman and Leopold 1987; Behrens *et al*. 1982; Iwabuchi *et al*. 1989). Similar pattern of H^+^ fluxes was determined in *Arabidopsis thaliana*, where the fluxes also correlated with root surface pH: an alkaline domain was observed at the border between meristematic and transition zones (Staal *et al*. 2011). Other works have attempted to determine the pH of the epidermal cell walls, and show a general decrease of pH towards the maturation zone of the root (Barbez *et al*. 2017; Moreau *et al*. 2022; Großeholz *et al*.2022). It is important to keep in mind that changes in proton fluxes and membrane potential are not always correlated with change in pH. For example, a simultaneous leak of protons and uptake through H^+^/K^+^ symporter in the cytoplasm can lead to apoplast acidification with a relatively stable membrane potential (Stanković 2006). In general, the relationship between H^+^ fluxes, cell wall pH and the growth rate in the particular zones remains unclear: the highest pH is observed in the transition and early elongation zones, while the lowest pH and highest outward H^+^ fluxes are observed in the maturation zone where cells do not elongate. In particular, the mechanism of the alkaline domain formation and its physiological significance remain unclear.

The apoplastic pH is largely controlled by the activity of the proton pump H^+^ATPases, encoded by 11 genes in the *Arabidopsis thaliana* genome; in roots, the dominant paralogs are AHA1 and AHA2 (Haruta *et al*. 2010). While their activity is mainly regulated by phosphorylation (Falhof *et al*. 2015), the expression and membrane localization of AHAs along the longitudinal root axis seems to be important for the zonation of the *Arabidopsis thaliana* root. Haruta *et al*. (2018) showed that under conditions of dim light, the AHA2-mCitrine was less abundant in the membranes of transition zone cells, and this partially correlated with higher surface pH of the transition zone cells. Under conditions of light, the AHA2-mCitrine localizes to the membranes in the transition zone. Pacifici *et al*. (2018) presented a model where cytokinin signaling regulates root meristem size by controlling the expression of AHA1 and AHA2. Increased cytokinin signaling led to shortening of the meristem, similarly to an inducible relocalization of AHA2 fused to the glucocorticoid receptor. Finally, Großeholz *et al*. (2022) quantified the AHA2-GFP signal and demonstrated an increasing AHA2-GFP membrane abundance towards the maturation zone of *Arabidopsis thaliana* root, which correlated with increased acidification of the root surface. Still, the observed patterns of proton fluxes along the root zones (Weisenseel and Meyer 1997), and in particular the existence of the alkaline domain around the transition zone of the root, cannot be simply explained by the expression or abundance of AHA proton pumps.

Roots use the gravity vector as the reference for the direction of their growth. A general pattern was shown in various plant species: during the gravitropic response, ion fluxes change rapidly, leading to increased proton secretion on the upper and decreased secretion on the lower side, where it correlates with growth inhibition, and results in bending of the root (Mulkey and Evans 1981; Behrens *et al*. 1982; Iwabuchi *et al*. 1989; Collings *et al*. 1992; Weisenseel *et al*. 1992). In *Arabidopsis thaliana*, the root surface and cell wall alkalinization of lower side upon gravistimulation has also been clearly demonstrated (Monshausen *et al*. 2011; Shih *et al*. 2015; Barbez *et al*. 2017), and it was shown that this alkalinization requires functional auxin transport (Monshausen *et al*. 2011) and the CNGC14 calcium channel acting downstream of auxin signaling (Shih *et al*. 2015). Alkalinization of the lower root side is thus triggered by the redirection of auxin flux towards the lateral root cap and epidermal cells on the lower side of the root (Brunoud *et al*. 2012). Analogously, apoplast alkalinization can be triggered by the application of auxin to the roots (Evans *et al*. 1980; Lüthen and Böttger 1988; Monshausen *et al*. 2011; Shih *et al*. 2015; Barbez *et al*. 2017). The root surface pH increases almost immediately upon auxin application, and this process requires the activity of the CNGC14 calcium channel; in the mutant, surface alkalinization as well as the auxin-induced growth inhibition are delayed (Shih *et al*. 2015). The rapid root growth inhibition as well as the membrane depolarization and surface alkalinization triggered by auxin depend on the rapid branch of the TIR1/AFB signaling branch; auxin influx by AUX1 and the AFB1 auxin co-receptor paralogue play a prominent role in the rapid auxin response (Fendrych *et al*. 2018; Dindas *et al*. 2018; Prigge *et al*. 2020; Serre *et al*. 2021; Li *et al*. 2021; Dubey *et al*. 2021). The molecular mechanism of surface alkalinization is, however, not understood. It was proposed that auxin plays a dual role in regulation of root apoplastic pH: it alkalinizes it by the activity of an unknown ion channel downstream of the TIR1/AFB signaling, and, at the same time, auxin upregulates the activity of AHA H^+^ ATPases through TMK1 signaling (Li *et al*. 2021). It is not clear how auxin activates the TMK kinases. On the other hand, the connection between TIR1/AFB auxin signaling and activity of AHAs has been clarified: the auxin-induced SAUR proteins inhibit the PP2C-D clade of protein phosphatases that act as negative regulators of AHAs, leading to the activation of proton secretion. This pathway was studied mainly in shoots, but seems also to operate in roots, as the Arabidopsis lines with manipulated SAUR or PP2C-D expression show root phenotypes that are consistent with the expected changes in AHA activities (Spartz *et al*. 2014; Ren *et al*. 2018). In summary, the connection between auxin signaling, regulation of the activity of AHA H^+^ ATPases and the longitudinal zonation of the root surface pH profile during normal growth and during gravitropic responses remains unclear.

Here, by employing an improved method for visualization and quantification of root surface pH, we focus on the molecular actors involved in the establishment of the longitudinal surface pH zonation. In particular, we show that the alkaline domain around the transition zone is not caused by the lack of AHA H^+^ ATPases activity. Further, we demonstrate that the alkaline domain at the transition zone is controlled by the components of the rapid auxin response pathway. Finally, we address the significance of the dynamic surface pH profile for the gravitropic response of the roots.

## Results

### The Arabidopsis root shows distinct acidic and alkaline root surface pH domains

To monitor the spatio-temporal dynamics of root ion fluxes and apoplastic pH in vertically growing roots of *Arabidopsis thaliana*, we re-evaluated the available fluorescence staining methods to visualize pH in roots. Cell wall staining by HPTS (Barbez *et al*. 2017) was not satisfactory in our setup, due to the very high background signal and probably due to the absence of the optimal 458 nm excitation in our vertical stage spinning disk microscope (Serre *et al*. 2021). However, at the transition zone of the root, we could observe an alkaline domain on the root surface (Fig.S1a). Further, we attempted to visualize root surface pH using Fluorescein Dextran and Oregon Green Dextran (Fig.S1a) pH reporters (Monshausen *et al*. 2011; Geilfus and Mühling 2011), but the results were not satisfactory due to an artifact when roots were imaged on solid medium (Fig.S1d). We therefore searched for alternative pH-sensitive fluorescent dyes, and discovered Fluorescein-5-(and-6)-Sulfonic Acid, Trisodium Salt (FS) (Invitrogen™ F1130; Seksek *et al*. 1995; Rosario and Rojas 1986) as an excellent reporter of root surface pH.

The F_488/405_ excitation ratio of FS efficiently reported pH when dissolved in liquid or solid medium (Fig.1a-d, Fig.S1 b,c). The F_488/405_ excitation ratio of FS is insensitive to the redox state of the medium but influenced by cation concentration (Fig.S1e). FS allows to visualize pH of the root surface and the surrounding rhizosphere, without penetrating the root tissues (Fig.1e). The pH imaging with FS highlighted the previously described alkaline domain in the transition zone/early elongation zone of the root (Monshausen *et al*. 2011; Staal *et al*. 2011). In addition, we observed two acidic pH domains, one located in the late elongation/root hair zone and the other in the proximity of the root tip (Fig.1e). To unbiasedly quantify the root surface pH, we developed a Python-based program to determine the F_488/405_ ratio of the root surface (Fig.S1f). The quantification highlighted the presence of the observed alkaline and acidic domains (Fig.1f).

**Figure 1:**
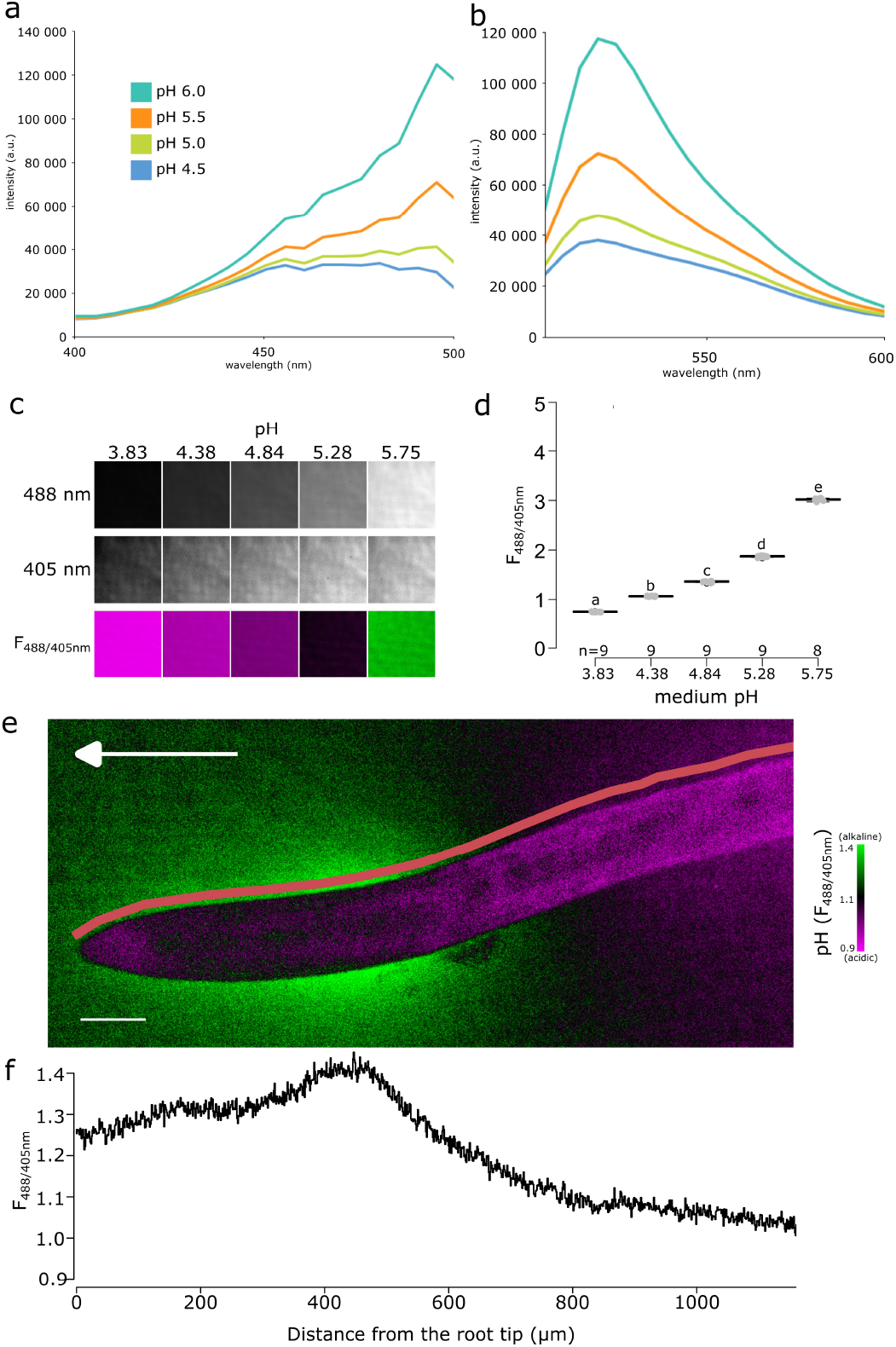
Fluorescein-5-(and-6)-Sulfonic Acid, Trisodium Salt dye reveals acidic and alkaline domains at the root root surface. (a,b) the pH dependence of the excitation (a) and emission (b) spectrum of Fluorescein-5-(and-6)-Sulfonic Acid, Trisodium Salt (FS) in liquid plant growth medium. Excitation spectra were recorded at λ_EM_ = 520 nm, emission spectra were excited by λ_EM_ = 488 nm. (c) FS fluorescence in solid agar medium at indicated pH values, fluorescence excited by 488 nm, 405 nm and the F_488/405_ excitation ratio, LUT as in (e). (d) Quantification of the F_488/405_ excitation ratio in (c). (e) Arabidopsis thaliana root tip shows the alkaline and acidic surface pH domains, arrow indicates the gravity vector, scale bar = 50 μm. The pink line shows the region in which F_488/405_ excitation ratio was plotted (f).

The surface pH profile of roots shown by FS is in agreement with the data obtained using numerous electrode and microscopy measurements in several species including *A. thaliana* (Zieschang *et al*. 1993; Staal *et al*. 2011; Monshausen *et al*. 1996; Weisenseel and Meyer 1997). The FS pH detection range thus reveals both the alkaline and acidic domains of the root surface and allows direct and dynamic visualization of proton fluxes between the root and its close surroundings.

### The alkaline domain in the transition zone is not directly determined by AHA activation or localization

We first hypothesized that the observed spatial surface pH gradients might originate from gradients of AHA ATPase abundance or activity (Großeholz *et al*. 2022). The immunolocalization of endogenous AHAs, however, didn’t reveal any obvious reduction in AHA abundance in the transition zone (Fig.2a, Fig.S2a) that would explain the presence of the alkaline domain. To overcome the hypothetical lack of AHA activation in the TZ, we treated seedlings with fusicoccin (FC), a fungal toxin that stimulates the AHA’s activity (Ballio *et al*. 1964) and thus increases proton efflux. FC lowered the surface pH of the root acidic domains but, surprisingly, did not affect the alkaline domain (Fig.2b,c). This partial acidification was correlated with the known FC-induced stimulation of root growth (Fig.S2b).

**Figure 2:**
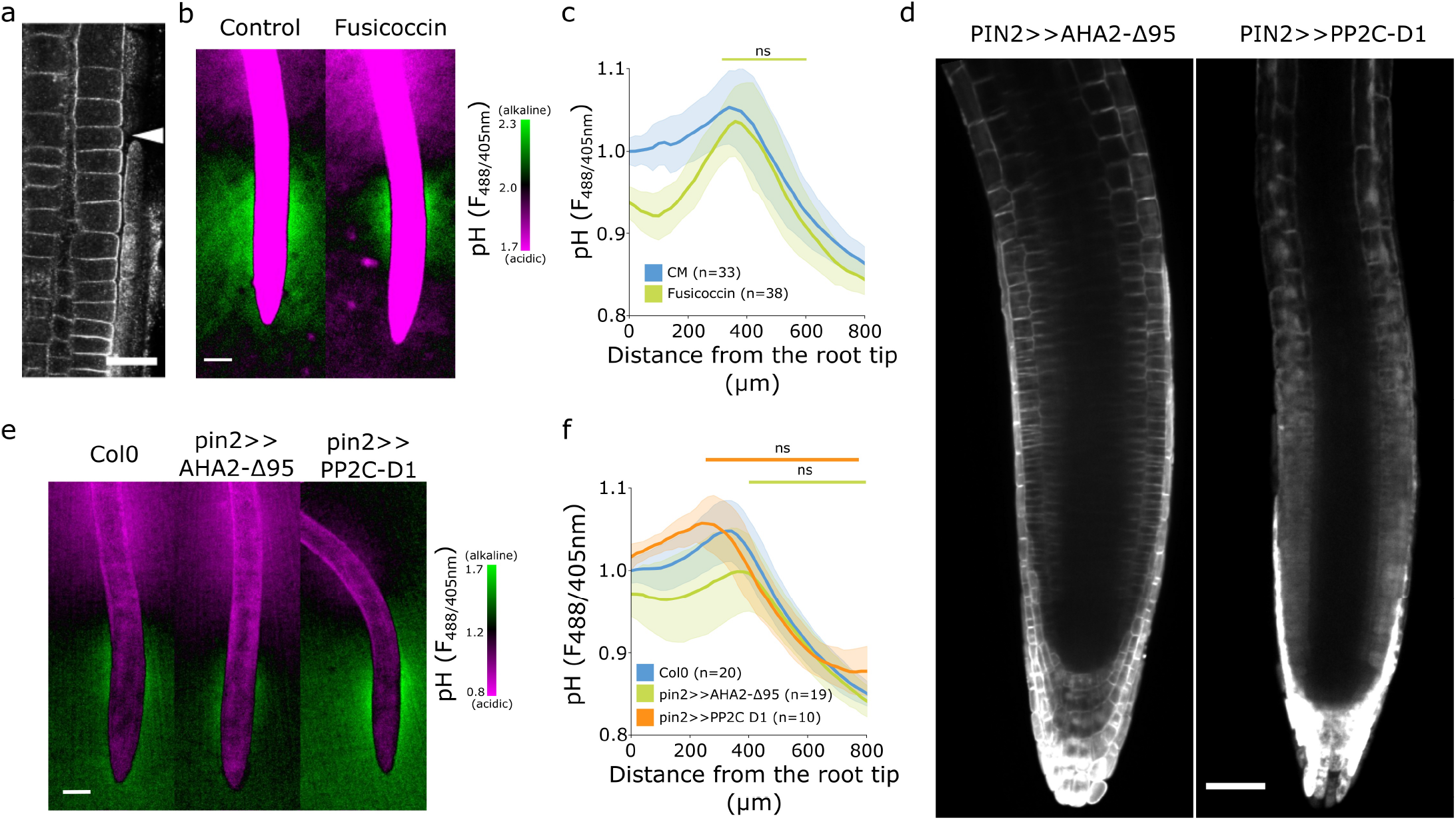
The alkaline domain does not directly depend on proton pump activity. (a) Immunostaining of AHA proton pumps in the Col-0 root transition zone (arrowhead). Scale bar = 20 μm. (b,c) Root surface pH visualized by FS of WT Col-0 seedlings treated with 0 μM (Control) or 2 μM Fusicoccin. (b) Representative image (scale bar = 100 μm) and (c) quantifications of FS F_488/405_ excitation ratio profile. (d) Image of root tips of the tissue-specific inducible lines PIN2>>AHA2-Δ95-mScarlet and PIN2>>PP2C-D1-mScarlet seedlings after 4-hour induction by 5 μM estradiol. Scale bar = 50 μm. (e,f) Root surface pH visualized by FS of WT Col-0 and induced PIN2>>AHA2-Δ95-mScarlet and PIN2>>PP2C-D1-mScarlet lines. (e) Representative images (scale bar = 50 μm), and (f) quantifications of FS F_488/405_ excitation ratio profile. For (c) and (f), ns = non significant statistical difference.

To exclude an FC-independent regulation of AHAs in the TZ, we created *Arabidopsis thaliana* lines inducibly expressing a fluorescently tagged hyperactive version of AHA2 (Pacheco-Villalobos *et al*. 2016) - AHA2-d95-mScarlet, and lines expressing an inhibitor of AHAs PP2CD1-mScarlet (Ren *et al*. 2018) under the control of the epidermal/cortex PIN2 promoter (Fig.2d). Expression of hyperactive AHA2 increased the acidic domain in the root tip, slightly reduced the alkaline domain, but did not prevent its formation (Fig.2e,f). This restricted acidification led to a tendency to stimulate root growth (Fig.S2c).

Inhibition of AHA activity by overexpression of PP2C-D1 raised the overall root tip surface pH, but did not prevent the formation of the alkaline domain in the TZ (Fig.2e,f). Alkalinization of the root tip surface correlated with a statistically insignificant reduction of root growth (Fig.S2c). We further measured the root surface pH profile of *aha2* and *pp2c-d triple* mutants that showed decreased (Haruta and Sussman 2012) and increased (Ren *et al*. 2018) AHA activity, respectively.

Plants lacking the expression of PP2CDs showed a significantly acidified root tip surface, however, the alkaline halo remained unaffected (Fig.S2g,i). On the other hand, plants lacking the expression of AHA2 were slightly impaired in the alkaline halo domain formation (Fig.S2g,i). *aha2-4* roots were growing slower than Col-0 while *pp2c-d triple* root growth was stimulated (Fig.S2h).

The manipulation of the AHA proton pump activity resulted in the expected outcome of influencing the overall root surface pH and root growth. However, the spatial organization of the root surface pH profile with the alkaline domain of the TZ does not appear to be simply controlled by the activation or abundance of AHAs, as the alkaline domain cannot be fully removed by genetic or pharmacological modulation of the proton pumping activity.

### The establishment of the alkaline pH domain requires AUX1

It is well established that the application of auxin to roots causes an increase in apoplastic and root surface pH (Monshausen *et al*. 2011; Shih *et al*. 2015; Li *et al*. 2021). To investigate the effect of auxin application on surface pH profile and formation of the TZ alkaline domain, we analyzed surface pH in response to the native auxin indole-3-acetic acid (IAA). With the increasing IAA concentration, the alkaline domain pH rose and the domain expanded towards the elongation zone of the root, which resulted in disappearance of the acidic domain (Fig.3a,b). The extent of the surface alkalinization correlated with IAA-induced root growth inhibition; the response was detectable at 1 nM IAA and was saturated between 100 and 1000 nM IAA (Fig.3c). This implies that the entire root surface is capable of alkalinization upon external auxin treatment, and that the alkaline domain is the hotspot of auxin response.

**Figure 3:**
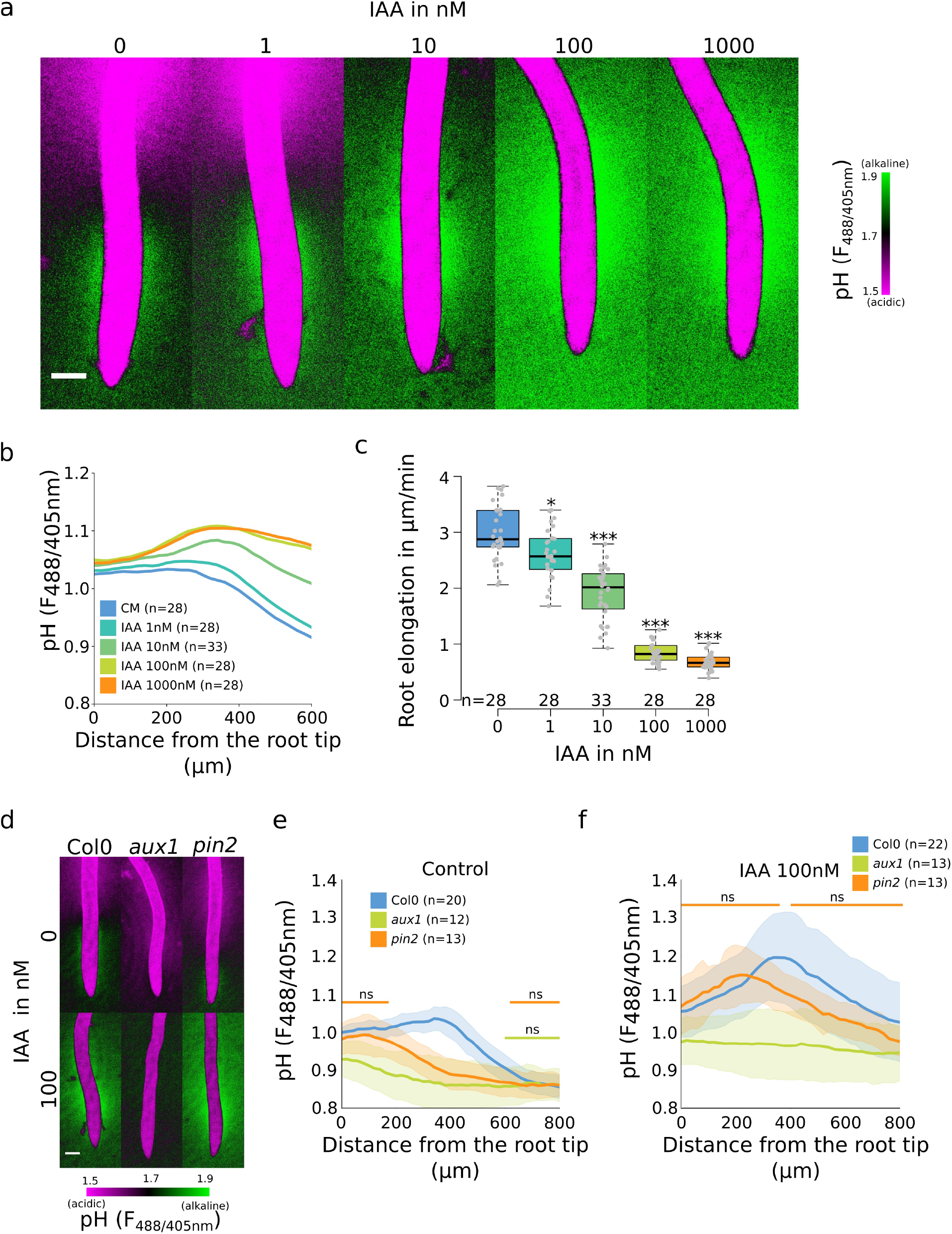
Auxin influx by AUX1 is essential for the initiation of the alkaline domain. (a-c) Surface pH correlates with growth rate - dose response of Col-0 roots to IAA auxin (a) Representative images of root surface pH visualized by FS after 20-min IAA treatment. (b) Quantifications of FS F_488/405_ excitation ratio profile. (c) Root elongation rate (μm/minute) measured over a 40 min period. (d-f) Root surface pH visualized by FS of Col-0, *aux1* and *pin2* mutants after 20 min treatment with 0 or 100 nM IAA. (d) Representative images. (e) Quantification of FS F_488/405_ excitation ratio profile in (e) control condition and (f) in response to 100 nM IAA. For b,e,f ns= non significant statistical difference. For c, *:p-value<0.05, ***:p-value<0.0005. Scale bars = 100 μm.

We further analyzed the root surface pH response to IAA in lines with altered AHA activity. The PIN2>>AHA2-d95 and PIN2>>PP2CD1 (Fig.S2d,e) as well as the *pp2c-d triple* mutant responded to IAA treatment by alkalinization of the root surface pH (Fig.S2i,j). The alkalinization factors (a measure of IAA-induced alkalinization) were similar to Col-0 with the exception of the very root tip of *pp2c-d triple* which responded slightly less (Fig.S2f,k). These results were correlated with control level IAA-induced root growth inhibition (Fig.S2c,h). On the other hand, knockout mutation of AHA2 led to reduced IAA-induced root surface alkalinization (Fig.S2i,j,k), but strong enough to observe an auxin-induced root growth inhibition (Fig.S2h).

The alkaline domain does not seem to be caused by the lack of proton efflux in the TZ, and the amplitude of the alkaline domain can be increased by auxin application. We therefore tested how auxin transport, perception and response contribute to the spatial determination of root surface pH.

First, we tested the mutant in the PIN2 auxin efflux carrier, in which the shootward auxin flux through the outer root tissues is perturbed (Luschnig *et al*. 1998; Müller *et al*. 1998). The *pin2* mutant root showed a reduced alkaline domain that was shifted towards the root tip (Fig.3d,e). Upon application of 10 nM IAA to *pin2* mutant, the alkaline domain position and amplitude were partially restored (Fig.S3a,b), the *pin2* roots also responded to auxin by growth inhibition (Fig.S3c). Upon treatment with 100 nM IAA, the alkaline domain and the overall IAA-induced root alkalinization were fully restored (Fig.3d,f and S3d,e).

The auxin influx carrier AUX1 is essential for auxin uptake and transport during root gravitropism (Swarup *et al*. 2005; Band *et al*. 2014), and the null *aux1* mutant was shown to have a more acidic surface with an altered root tip pH profile (Monshausen *et al*. 2011). We determined the root surface pH profile in the *aux1* mutant, and found that its pH showed a gradual decrease from the tip towards the root hair zone. While the acidic zone in the elongation zone was comparable to control roots, *aux1* mutants displayed a complete absence of the TZ alkaline domain (Fig.3d,e). Application of 10 nM IAA on *aux1* mutants did not affect root elongation (Fig.S3c) and did not trigger a root surface alkalinization in contrast to the control Col-0 (Fig.S3a,b). However, at 100 nM IAA, when IAA diffusion compensates for AUX1 function (Evans *et al*. 1994), *aux1* mutant showed a clear IAA-induced root growth inhibition (Fig.S3e) as well as an increased root surface pH beyond the TZ (Fig.3d,f and S3d). The WT-like root surface pH profile was, however, not restored by IAA application, demonstrating that AUX1 is essential for creating the alkaline surface domain in the transition zone.

As the immunolocalization of AHAs in the *aux1* mutant was comparable to the Col-0 control (FigS2a), we treated the mutant with FC to investigate the activation status of AHAs in the *aux1* mutant. Interestingly, this treatment resulted in the establishment of a small alkaline domain also in the *aux1* mutant (Fig.S3f,h), likely caused by AHA-mediated acidification in the root tip and the distal elongation zone that did not affect the TZ surface pH. The partial acidification was again correlated with a significant root growth stimulation (Fig.S3g). This experiment showed that AHAs are not fully activated in *aux1* mutant and implies a specific surface pH response of the cells of the TZ.

In summary, we have shown that the distinctive alkaline domain at the transition zone of the root depends on AUX1-mediated auxin influx. Without auxin influx, the root shows a linear acidification gradient from the root tip to the maturation zone.

### The components of the rapid auxin response pathway steer the transition zone root surface pH

Once in the cytoplasm, auxin triggers responses either through the canonical auxin signaling pathway or the auxin rapid response pathway (Dubey *et al*. 2021). We examined the involvement of these pathways in the establishment of the surface pH profile of the root. First, we investigated the components of the canonical auxin signaling by using the *tir1,afb2,3 (tir triple)* mutant which lacks the expression of three of the six known auxin receptors (Dharmasiri *et al*. 2005). The *tir triple* mutant displayed a more acidic surface pH, particularly in the elongation and maturation zone (Fig.4a,b). However, it was still displaying an alkaline domain, albeit less pronounced than the control. In response to IAA, *tir triple* was critically impaired in whole root IAA-induced surface pH alkalinization (Fig.S4b,c). As a result, *tir triple* was also impaired in the IAA induced root growth inhibition (Fig.S4a).

**Figure 4:**
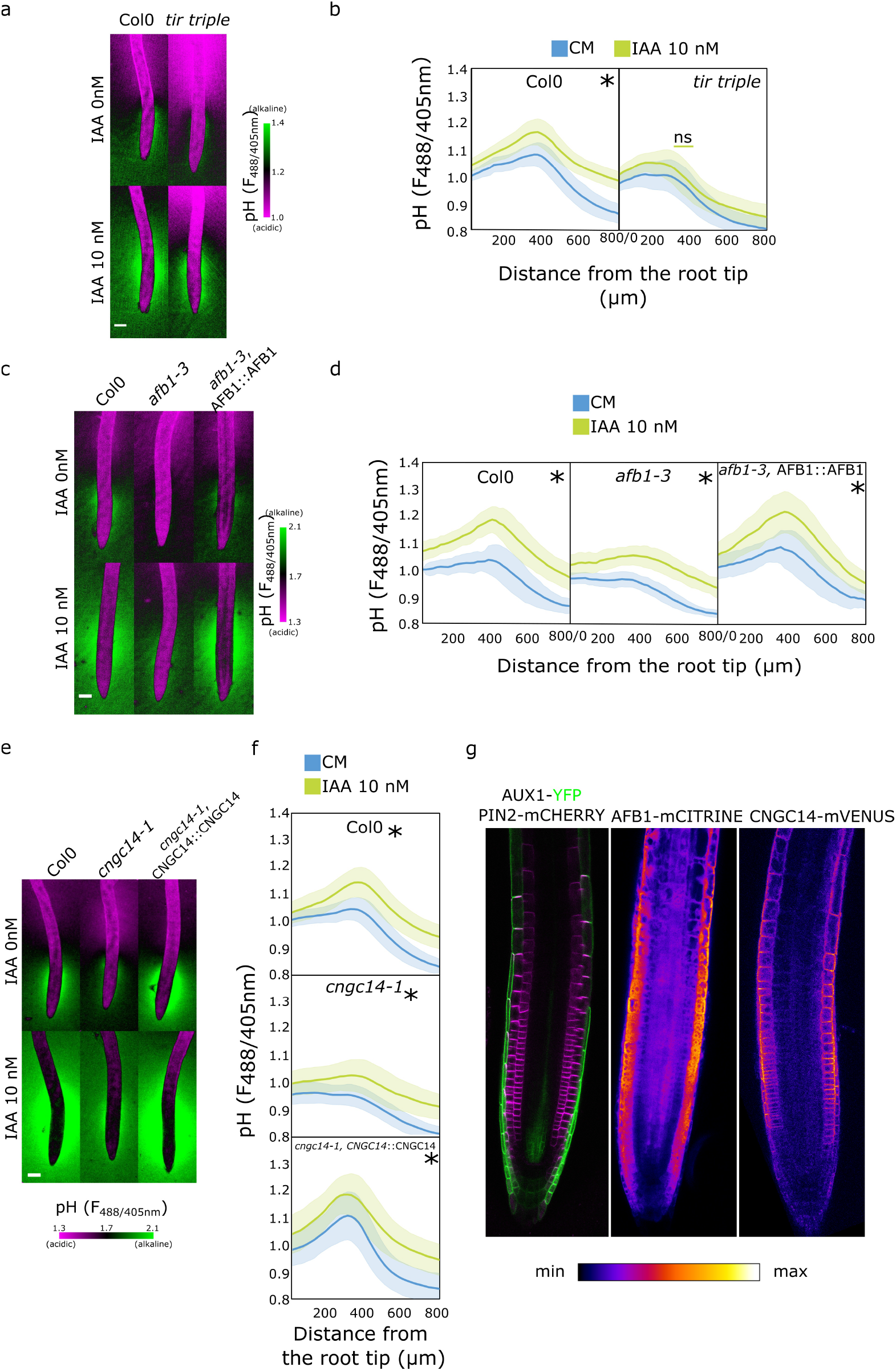
Rapid auxin signaling steers root surface pH. (a, b) Root surface pH of Col-0 and *tir triple* mutant after 20-min treatment with 0 or 10 nM IAA. (a) Representative images of pH visualization by FS and (b) quantification of FS F_488/405_ excitation ratio profile. (c-d) Root surface pH of Col-0, *afb1-3* mutant and AFB1::AFB1-mCitrine/*afb1-3* complemented line after 20-min treatment with 0 or 10 nM IAA. (c) Representative images of pH visualization by FS and (d) quantification of FS F_488/405_ excitation ratio.(e-f) Root surface pH of Col-0, *cngc14-1* mutant and CNGC14::CNGC14-mVenus/*cngc14-1* complemented line after 20-min treatment with 0 or 10 nM IAA. (e) Representative images of pH visualization by FS and (f) quantification of FS F_488/405_ excitation ratio. (g) Localization of AUX1, PIN2, AFB1 and CNGC14 proteins driven by their respective native promoters. For b,d,f, ns = non significant statistical difference. Scale bars = 100 μm (a,c,e) or 50 μm (g).

These results confirm that the canonical auxin signaling is involved in the auxin-induced apoplastic alkalinization (Li *et al*. 2021) and also partially in the longitudinal surface pH zonation.

We further explored the role of the recently discovered molecular actors of the auxin rapid response - AFB1 and CNGC14. AFB1 is the paralogue of the TIR1 receptor; AFB1 has been shown to be crucial for the rapid growth inhibition upon auxin treatment, rapid auxin-induced membrane depolarization, and for the early response to gravistimulation (Prigge *et al*. 2020; Serre *et al*. 2021). Similarly, the mutant in the calcium channel CNGC14 lacks the auxin-induced calcium transient, membrane depolarization, and shows a delay in the early gravitropic response (Shih *et al*. 2015; Dindas *et al*. 2018). We analyzed the root surface pH of both mutants and found a more acidic root surface with a critically flat alkaline domain in the *afb1* mutant (Fig.4c,d and S4d) as well as in the *cngc14* mutant (Fig.4e,f and S4e). We confirmed the similar result in additional alleles of *afb1* (Fig.S4f) and *cngc14* mutants (Fig.S4g). This shows that these proteins, apart from controlling the rapid auxin response, steer the pH profile of the root during normal steady state growth by controlling the alkaline domain in the transition zone. The lack of the alkaline domain was not caused by mislocalization or absence of AUX1 protein in the *afb1* or *cngc14* mutants (Fig.S4h). Similarly to the *aux1* mutant, the localization of AHA H^+^ ATPases in both mutants was similar to the control (Fig.S2a). We further tested how the *afb1* and *cngc14* mutants respond to IAA and found an overall alkalinization of the root surface with a partial rescue of the alkaline domain (Fig.4d,f) and partially impaired growth inhibition (Fig.S4i-k).

What makes the TZ zone surface pH different from the other domains of the roots? A plausible explanation is the localization of relevant molecular components in this region of the root. The auxin transporters AUX1 and PIN2 have been shown to localize to the lateral root cap and epidermis (Müller *et al*. 1998; Swarup *et al*. 2004, Fig.4g). Prigge *et al*. (2020) showed that AFB1 is expressed in the root tip and we could see an enrichment of the protein in the root epidermis (Fig.4g). To determine the localization of CNGC14, we expressed its fluorescently C-terminally tagged version under the control of its native promoter.

The expression of CNGC14 fusion protein was rather weak, and localized to the plasma membrane of root epidermal cells, with an enrichment in the transition zone (Fig.4g). The expression of AFB1-mCitrine and CNGC14-mVenus in the respective mutants recovered the presence of the TZ alkaline domain (Fig.4c,d,e,f and S4d,e,g). Interestingly, the expression of CNGC14-mVenus in *cngc14* caused an exaggerated TZ alkaline domain (Fig.4e,f and S4e,g), which underlines the importance of the CNGC14 calcium channel in the establishment of root surface pH profile.

These results show that the signaling components so far associated with the rapid auxin response are expressed in the root epidermis and contribute to the longitudinal zonation of root surface pH profile. The alkaline domain of the TZ represents a site of a constant rapid auxin response that is triggered by the internal auxin fluxes mediated by AUX1 and PIN2 transporters. On the other hand, the overall increase of root surface pH upon auxin treatment can be attributed to the TIR1 canonical signaling pathway.

### The AUX1-AFB1-CNGC14 module facilitates root navigation

It was shown that auxin induces alkalinization of the apoplastic pH and that this alkalinization correlates and is required for the inhibition of root growth. The same process occurs during the gravitropic response - the lower root side responds to the internal auxin by alkalinization, resulting in root bending (Monshausen *et al*. 2011; Shih *et al*. 2015; Barbez *et al*. 2017). We reanalyzed the root surface pH using our pH imaging and quantifications, as it enables us to monitor alkalinization, as well as acidification of the root surface. Upon gravistimulation, the TZ alkaline domain on the lower side of the root increased rapidly, while on the upper side, the alkaline domain diminished and the root surface acidified (Fig.5a, Supplemental Movie 1). This led to a gradient of surface pH across the root that was established within 5 minutes of the gravistimulation which correlated with initiation of bending of the root (Fig.5a). The disappearance of the alkaline halo on the upper root surface indicates that the TZ domain might act as a zone of stalled growth which is rapidly activated upon gravistimulation. We further analyzed the gravitropic responses of the mutants with altered root surface pH zonation. As expected, the agravitropic *aux1* mutant did not create a pH gradient across the root and did not bend (Fig.5b). Manipulation of the AHA activity by the application of fusicoccin or by genetic means did not prevent the formation of a gradient of surface pH and rapid bending of the root (Fig.S5a,b). Finally, the *afb1* and *cngc14* mutants showed a slower gravitropic response, as reported before (Serre *et al*. 2021; Shih *et al*. 2015) (Fig.5c,d). As both mutants have a diminished alkaline domain in the TZ, upon gravistimulation, this zone could not rapidly react to change in auxin fluxes; instead, a shallow gradient of surface pH slowly develops in the mutants (Fig.5 c,d).

**Figure 5:**
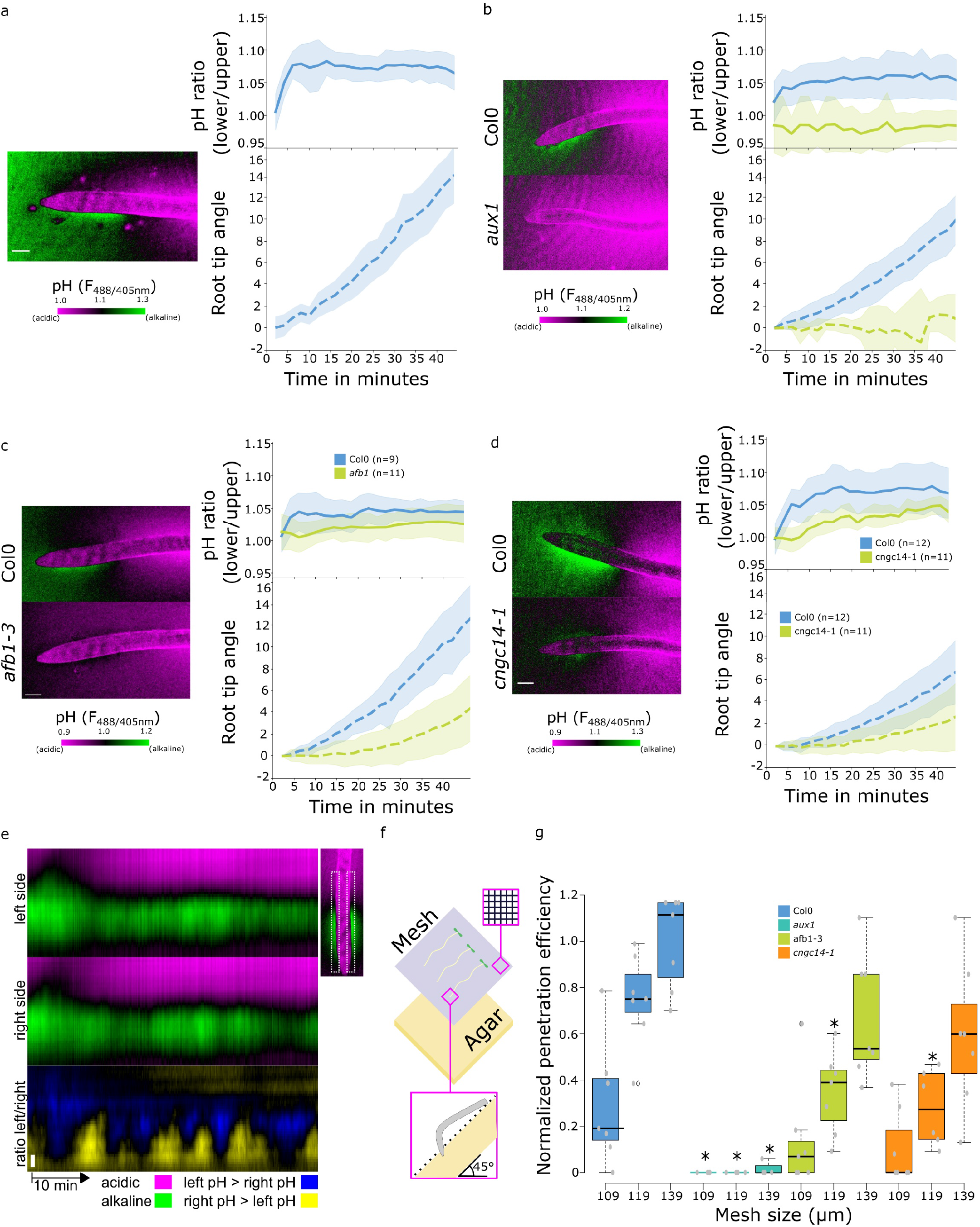
Rapid auxin signaling pathway is required for surface alkalinization during rapid gravitropic responses. (a-c) Surface pH dynamics and root tip bending angle during gravitropic response in (a) Col-0, (b) Col-0 and *aux1*, (c) Col-0 and *afb1-3*, (d) Col-0 and *cngc14-1* lines. A representative image of pH visualization by FS, quantification of the FS F_488/405_ excitation ratio of lower/upper root transition zones, and root tip angle dynamics over time are shown for each line. Representative images were taken 40 minutes after gravistimulation. (e) Root surface pH oscillations in vertically growing Col-0 roots. The FS F_488/405_ excitation ratio images for left and right root sides and their left/right ratio are shown. (f,g) Root tip penetration test in 45° inclined media covered with mesh of different pore sizes (109, 119 and 139 μm). (f) Schematics of the experimental setup. (g) Quantitation of Col-0, *aux1, afb1-3* and *cngc14-1* mesh penetration efficiencies. For g, statistical differences with p-value<0.05 indicated by *. All scale bars = 100 μm

We noticed that the alkaline domain showed dynamic fluctuations during vertical growth of the root (Fig.5e, Supplemental movie 2), similarly to what was described by Monshausen *et al*. (2011). This result and the fact that the components of the rapid auxin response pathway are involved in the formation of the alkaline domain indicates that, in this zone, the cells continuously rapidly respond to the internal auxin fluxes, and that this pathway is active not only upon gravistimulation. We hypothesized that the significance of this process is to constantly correct the growth rate fluctuations and to enable the root to quickly regulate the growth direction. To test this hypothesis, we tested the ability of WT and *aux1*, *afb1* and *cngc14* mutants to navigate through artificially created obstacles. We scored the efficiency of the roots to penetrate tilted nylon grids with varying pore sizes (Fig.5f). WT plants penetrated the meshes more efficiently as the pore size increased, whereas the agravitropic *aux1* mutant failed to penetrate in all conditions. Interestingly, both the *afb1* and *cngc14* mutants showed a significant decrease in penetration efficiency compared to the wild type (Fig.5g).

In summary, the alkaline surface pH domain originates from a constant rapid auxin response driven by the AUX1 - AFB1 - CNGC14 module. This response enables the roots to rapidly react to gravity vector changes and to adjust the root tip growth direction.

## Discussion

In this work, we present a new method to determine the root surface pH of *Arabidopsis thaliana*. Its advantage is the sensitivity of FS in lower pH range that enables a spectacular visualization of root surface pH, including the acidic domain surrounding the late elongation and maturation zones of roots, which was not detected with the previously published methods (Monshausen *et al*. 2011; Shih *et al*. 2015). FS is an inexpensive dye which can be excited by the common 405 and 488 nm laser lines. In this work, we avoided measuring absolute pH values; instead, we determined the relative pH compared to internal controls observed on the same medium. Absolute pH determination would be possible, if extensive pH calibration was performed during each imaging session. This approach would be very laborious, and, in addition, absolute fluorescence intensities might depend on the thickness of the layer of medium and many other parameters. Therefore, we recommend using FS to determine relative pH changes compared to internal controls. We showed that apart from pH, FS can also react to other ions, however, given that the surface pH profile visualized by FS staining is in full accordance with the previously published works (Weisenseel and Meyer 1997; Weisenseel *et al*. 1979; Björkman and Leopold 1987; Behrens *et al*. 1982; Iwabuchi *et al*. 1989; Staal *et al*. 2011), we conclude that in the growth media, FS reflects primarily the root surface pH.

According to the acid growth theory, acidic pH enables cell wall extension (Rayle and Cleland 1992; Hager 2003). We observed the lowest pH values in the very tip of the root and in the maturation zone where cells do not elongate. On the other hand, the alkaline domain partially covers the early elongation zone. The surface pH therefore does not fully correlate with the local growth rates (Beemster and Baskin 1998; Shih *et al*. 2015). In addition, it remains to be resolved to what extent the surface pH correlates with the pH of the cell walls and the apoplastic space of the root epidermal cells (Barbez *et al*. 2017; Moreau *et al*. 2022).

It is possible that rather than driving cell elongation, the acidic domain in the maturation zone plays a role in nutrient acquisition and absorption by the root hairs (Martín-Barranco *et al*. 2021). The cytokinin-mediated cell wall stiffening may become the dominant growth-regulating mechanism in this root zone, leading to growth cessation despite low extracellular pH (Liu *et al*. 2022).

Based on genetic and pharmacological experiments we concluded that the longitudinal surface pH zonation is not based solely on the activity and abundance of PM AHA H^+^ATPases, but instead, the alkaline domain in the transition zone is driven by AUX1-mediated auxin influx and AFB1 signaling. The surface pH might be determined solely by AHA activity in the specific case of the *aux1* mutant, where the pH progressively decreases in the shootward direction, and might then reflect the AHA activity gradient that is determined by the abundance of the proton pumps and their regulation by brassinosteroid signaling, as suggested by Großeholz *et al*. (2022). Interestingly, even in the *aux1* mutant, a residual AHA-independent alkaline domain is present and can be visualized when AHAs are hyperactivated by the application of fusicoccin.

In agreement with these results, the ectopic AHA ATPases activation also cannot prevent the auxin-induced growth inhibition and the gravitropic bending of roots. Inhibition of AHA activity led to slower growth, a tendency of slower gravitropic response, and the *aha2-4* mutation caused a reduced auxin-induced surface alkalinization. In addition to controlling membrane potential and apoplastic pH, AHA activity influences auxin signaling by regulating auxin influx and diffusion into cells (Rubery and Sheldrake 1973; Yang *et al*. 2006). These results might thus be caused by the reduced proton motive force that in turn might interfere with IAA uptake into the cells.

What is the cause of the alkaline domain at the transition zone, if not the inhibition of AHA ATPase activity? It was shown that the auxin-induced apoplast and root surface alkalinization is accompanied by a decrease in cytosolic pH (Monshausen *et al*. 2011; Li *et al*. 2021), indicating that the alkaline domain visualized by the FS is caused by a proton influx into cells. The phenotype of the *aux1* mutant hints to the possibility that the AUX1-mediated auxin symport with protons (Lomax *et al*. 1985) might cause the observed root surface alkalinization. Also supporting this hypothesis, the transition zone of maize roots is the site of the highest auxin influx, as measured by an auxin-specific electrode (Mancuso *et al*. 2005). On the other hand, the *aux1* mutant was capable of surface alkalinization when higher concentrations of IAA were added, demonstrating that the AUX1-mediated influx *per se* is not required for surface alkalinization. Further, the *afb1* mutants show a defect in the alkaline domain formation, which favors the explanation that the auxin-induced alkalinization is triggered from the AFB1 receptor. Given that the mutant in CNGC14 is defective in establishing root surface pH profile, auxin-induced alkalinization and auxin-induced calcium influx (Shih *et al*. 2015), and calcium influx and pH changes are tightly coupled (Behera *et al*. 2018), we conclude that CNGC14 functions downstream of AFB1. Similarly, Li *et al*. (2021) suggested that auxin-induced alkalinization of the apoplast is mediated by a yet unknown mechanism. It is intriguing to speculate that the signaling pathway might operate via the adenylate cyclase activity of TIR1/AFB auxin receptors (Qi *et al*. 2022).

We show that the rapid auxin response pathway, so far connected with the reaction to auxin application (Fendrych *et al*. 2018; Li *et al*. 2021; Monshausen *et al*. 2011) or the gravitropic response (Serre *et al*. 2021; Shih *et al*. 2015), operates constantly in the growing *Arabidopsis thaliana* root. Even though the other TIR1/AFB coreceptors partially contribute to the longitudinal surface pH zonation, AFB1 plays the most important role. The other TIR1/AFB receptors seem to be more important for the overall pH rise upon the application of IAA, in agreement with the results of Li *et al*. (2021). The localization of the alkaline zone seems to be determined by the intersection of the PIN2-mediated shootward auxin flux (Luschnig *et al*. 1998; Müller *et al*. 1998), localization of the AUX1-mediated auxin influx (Swarup *et al*. 2005; Band *et al*. 2014) and the enrichment of the AFB1 and CNGC14 proteins. We propose that the dynamic nature of the surface alkalinization observed by us and others (Monshausen *et al*. 2011; Shih *et al*. 2015) results from the interaction of the intensity of the auxin flux and the constant rapid response that occurs in the alkaline domain of the root tip. In addition, the auxin-induced alkalinization is constantly being counterbalanced by the TMK-mediated AHA ATPase activation and apoplast acidification (Li *et al*. 2021; Friml *et al*. 2022; Lin *et al*. 2021).

Upon gravistimulation, the alkaline domain on the lower side of the root increased rapidly, while on the upper side, the alkaline domain diminished and the root surface acidified, which is consistent with previous pH measurements done by electrodes (Zieschang *et al*. 1993; Monshausen *et al*. 1996), indicating that the alkaline domain corresponds to a zone of a stalled cell elongation that can be rapidly regulated upon gravistimulation. In addition to the gene expression changes that occur during obstacle avoidance (Jacobsen *et al*. 2021) and the FERONIA-mediated mechanoperception (Shih *et al*. 2014), the rapid auxin response module in the root decreases the reaction time of the root to the changes of root growth direction, and thus increases the efficiency of root soil penetration.

## Material and methods

### Plant material used

We used the Col-0 ecotype and the following mutant and transgenic lines. The PIN2>>AHA2delta95, PIN2>>PP2C-D1 originate from this study, see Molecular cloning. Further, we used the *aha2-4* (SALK_082786), *pp2c-d2/5/6* triple mutant (Ren *et al*. 2018) (pp2c-d2 WsDsLOX493G12, pp2c-d5GABI_330E08 and pp2c-d6 SAIL_171H03), *aux1-100* (SALK_020355), *pin2* (NASC_N16706). The *afb1-3* and AFB1::AFB1-mCitrine in *afb1-3* originate from (Prigge *et al*.2020). The *afb1-1s* (SALK_144884C) was genotyped using the following primers SALK_LBb1.3, AACGGAAGACTAGGAAGCGAG, GCAACAGCTTCAAGACCTTTG. The *cngc14-1* (SALK_206460) was genotyped by SALK_LBb1.3, CACCTGCTTGTAAAGCAAAGG, TCGGAACAATTGGCAGAATAC; *cngc14-2* (wiscDsLox437E09) by WisDsLox_LBP (AACGTCCGCAATGTGTTATTAAGTTGTC), TGTTTCACGTAAAGTCAAACCC, TAAGAATCCAAGTGGCCACAC. The CNGC14::CNGC14-mVenus (see Molecular cloning) was transformed into the *cngc14-1* and *cngc14-2* homozygous lines. The *tir triple* is the *tir1,afb2,3* mutant (Dharmasiri *et al*. 2005). AUX1::AUX1-YFP (Swarup *et al*. 2004) was introduced into *afb1-3* and *cngc14-2* by crossing. AUX1::AUX1-YFP was crossed with PIN::PIN2-mCherry (Retzer *et al*. 2019).

Seeds were surface-sterilized by chlorine gas for 2h. Seeds were sown on 1% (w/v) plant agar (Duchefa) with ½ Murashige Skoog salts (MS, Duchefa), 1% (w/v) sucrose, adjusted to pH 5.8 with KOH, and stratified for 2 days at 4 °C. Seedlings were vertically grown for 5 days in a growth chamber at 23 °C by day (16h), 18 °C by night (8h), 60% humidity and light intensity of 120 μmol photons m-2 s-1.

### Molecular cloning

The tissue-specific estradiol-inducible lines were prepared as follows. For PIN2>>AHA2delta95, we cloned the CDS 1-886 of AHA2 (AT4G30190) lacking the last 95 amino acids as in (Pacheco-Villalobos *et al*. 2016), and fused mScarlet-I (Bindels *et al*. 2017) to the C-terminal part. For PIN2>>PP2C-D1, we cloned the CDS of PP2C-D1 (AT5G02760) and fused mScarlet-I to the C-terminal part. Both constructs were cloned downstream of the 4xLexA Operon fused to CaMV 35s minimal promoter (Sarrion-Perdigones *et al*. 2013) and the transcriptional units were terminated by the 35S terminator and cloned into the alpha1-3 vectors (Dusek *et al*. 2020). XVE (Zuo *et al*. 2000) was cloned under the control of the PIN2 promoter (1.4 kb upstream of the AT5G57090), terminated by the RuBisCo terminator from *Pisum sativum* and the resulting transcriptional unit was cloned into alpha1-1 vector. The alpha transcriptional units were then interspaced with matrix attachment regions (Dusek *et al*. 2020), combined with a Basta resistance cassette and introduced into the pDGB3omega1 binary vector (Sarrion-Perdigones *et al*. 2013). The CNGC14::CNGC14-mVenus construct contains the CNCG14 promoter (1.5 kb upstream of the start codon) and the CNGC14 coding sequence (the AT2G24610.1 splice variant) with the mVenus fused to the C-terminus and 35S terminator. This transcriptional unit was combined with a kanamycin resistance cassette and FastRed marker for rapid selection of transgenic seeds into the pDGB3omega1 binary vector. All cloning steps were performed using the GoldenBraid methodology (Sarrion-Perdigones *et al*. 2013; https://gbcloning.upv.es/).

Col-0 ecotype (PIN2>>AHA2delta95, PIN2>>PP2C-D1) or *cngc14-1* and *cngc14-2* lines (CNGC14-mVenus) were transformed using the floral dip method (Clough and Bent 1998).

### Pharmacological treatments and dyes

Treatments were prepared using the following chemicals: IAA (10 mM stock in 70 % ethanol; Sigma Aldrich), fusicoccin (1 mM stock in 70 % ethanol; Sigma Aldrich), estradiol (20 mM stock in DMSO; Sigma Aldrich). Fluorescein Dextran 10 000 MW, Anionic (D1821, ThermoFisher)-10 mg/ml stock in miliQ H2O, final concentration in media is 29 μg/ml. Oregon Green 488 Dextran, 10 000 MW (D7170, Invitrogen) −1 mg/ml stock in miliQ H2O, final concentration in media is 2.5 μg/ml. HPTS −8-Hydroxypyrene-1,3,6-trisulfonic acid trisodium salt (H1529, Sigma Aldrich), 100 mM stock in H2O, final concentration in media is 1 mM. FS - Fluorescein-5-(and-6)-Sulfonic Acid, Trisodium Salt (F1130, ThermoFisher), 100 or 50 mM stock in H2O, final concentration in media is 50 μM.

Induction of PIN2>>AHA2delta95 and PIN2>>PP2C-D1 lines was done by incubating 5-day-old seedlings for 2.5 hours in 1/2MS MES buffered media (g/L), 1 % sucrose, pH 5.8 containing 2 μM estradiol before experiments.

### Measurement of FS excitation and emission spectra

For measurement of fluorescent spectra of FS, ½ MS media with 1 % sucrose were prepared with pH 4.5, 5.0, 5.5, 6.0 (adjusted with KOH). FS stock (50 mM in H2O) was added to the media to achieve a final concentration of 50 μM. For the measurement, 200 μl of this media was pipetted into a 96-well plate, five wells per each pH value and the spectras were measured using Spark® multimode microplate reader (Tecan). The values for each pH were averaged and plotted.

### Measurement of FS range and effects of salts and redox status on FS signal

To demonstrate the wide pH-reporting range of FS (Fig 1c), citric acid and sodium citrate were mixed in various ratios and water was added to obtain citrate buffers with a range of pH values. The pH of citrate buffers was measured, the buffers were mixed with agar (0.8% final concentration), boiled and FS was added (50 μM final concentration). The mixture was poured into a Petri dish. After cooling down, slabs of the solidified buffers were cut out, and placed into imaging chambers.

To measure the effects of ions and redox status (according to Martinière *et al*. 2013) on FS fluorescence (FigS1e), MES buffer (1g/l) was prepared and boiled with agar (1 % final concentration), supplied with salts, H2O2, DTT and FS was added (50 μM final concentration). The mixture was poured into a Petri dish. After cooling down, slabs of the solidified buffers were cut out, placed into imaging chambers and imaged using a spinning disk microscope, see *Imaging of root surface pH*. The signal intensity was measured in both 405 nm and 488 nm excitation channels and F_488/405_ ratio was calculated.

### Imaging of root surface pH

For imaging of surface pH, 5-day-old seedlings were transferred to unbuffered ½ MS, 1 % sucrose, pH 5.7 (adjusted with KOH) containing 50 μM of FS +/−treatments and allowed for recovery 25 minutes before imaging. The medium was prepared as follows: For 100 ml of medium, 1 g of plant agar and 1 g sucrose (w/v) were added to 100 ml of unbuffered ½ MS in a 250 ml reagent bottle. The media was boiled until the agar was completely dissolved. Note: we always prepared a fresh solid medium for each experiment from a stock of liquid ½ MS, sucrose and agar, as re-boiling the medium might affect the results. The medium was cooled to a temperature of 45-50 °C and FS was added to achieve 50 μM concentration. This pre-solution was then divided into two equal volumes to make control and treatment medium.

For the gravitropic experiments, the seedlings were rotated +/−90°. Only roots with a starting angle of 90°+/−10° were selected for analysis to obtain homogeneous gravitropic stimulations.

Surface pH imaging was performed using a vertical stage Zeiss Axio Observer 7 with Zeiss Plan-Apochromat 10×/0.8, coupled to a Yokogawa CSU-W1-T2 spinning disk unit with 50 μm pinholes and equipped with a VS401 HOM1000 excitation light homogenizer (Visitron Systems). Images were acquired using the VisiView software (Visitron Systems).

FS was sequentially excited with a 488 nm and 405 nm laser and the emission was filtered by a 500–550 nm bandpass filter. Signal was detected using a PRIME-95B Back-Illuminated sCMOS Camera (1200 × 1200 pixels; Photometrics). The Flat field correction mode was used for image acquisition.

For vertical growth experiments, seedlings were imaged every 10 minutes for 30 minutes. The profiles shown correspond to the first time frame while the root elongation was calculated over the whole experiment. Imaging to observe oscillations of the root surface pH were conducted by imaging every 5 seconds.

For gravitropic experiments, seedlings were imaged using the sandwich method (https://doi.org/10.1017/qpb.2022.4) every 2 minutes for 42 minutes. The profile shown corresponds to the last time frame except specified otherwise. The angles were quantified over the entire duration of the experiment.

The images in Fig.4g were acquired using the Zeiss 880 confocal microscope, with a 25×/0.8 water immersion objective using settings appropriate for the respective fluorophores.

### Immunolabeling of AHAs

Whole mount immunolocalization of 5–day–old Arabidopsis thaliana seedlings was performed as described previously in(Sauer *et al*. 2006), with the following modifications. The protocol was adapted to the InSituPro VS liquid-handling robot (Intavis AG, Germany). Prior to immunolocalization seedlings were fixed 1 h with 4% paraformaldehyde dissolved in MTSB (50 mM PIPES, 5 mM EGTA, 5 mM MgSO4·7H2O pH 7, adjusted with KOH) at room temperature. In the robot, procedure started with 5×15min washes with MTSB-T (MTSB+0.01% TritonX-100) then the cell wall was digested by 30 min treatment at 37°C with 0,05% Pectolyase Y-23 supplemented with 0,4M mannitol in MTSB-T, followed by 2×30 min membrane permeation with 10% DMSO and 3% Igepal in MTSB-T. The samples were blocked 1h with BSA (blocking solution: 2% BSA in MTSB-T) and incubated 4 h at 37°C with primary (anti-AHA rabbit antibody, Agrisera AS07260) and 3 h at 37°C with secondary antibody (Alexafluor 555 goat, anti-rabbit, Abcam ab150078). The antibodies were diluted in 2% BSA in MTSB-T in concentration: 1:500 for primary and 1:1000 for secondary antibody. Between the all described steps the seedlings were washed 5×15 min with MTSB-T. For the final step, MTSB-T was exchanged by deionized water. From the robot seedlings were transferred to microscopy slides into 50-% glycerol in deionized water and fluorescence signal was imaged by Zeiss LSM 880 inverted confocal scanning microscope equipped with Airyscan detector with 40×/1.2 C-Apochromat objective.

### Mesh penetration test

Four-day-old seedlings were transferred on ½ MS, 1% (w/v) plant agar, 1% (w/v) sucrose, pH 5.8, covered with sterilized nylon meshes of pore sizes of 109, 119, 139 μm (polyamide mesh UHELON). Plates were grown overnight vertically and then were tilted to 45 degrees and grown 24 hours. After that, the root penetration through the mesh was scored using a binocular microscope. The penetration efficiency was calculated by normalizing each variant (genotype X pore size) to the Col-0 efficiency in the 139 μm variant in the particular repetition. In addition, the reference Col-0 was normalized to the Col-0 average over all repeats.

### Image Analysis

We determined root elongation by measuring the total root length increment between the first and last time frame divided by the number of minutes. The root length increment was measured with the segmented line in ImageJ/Fiji v1.53f51 (Schindelin *et al*. 2012).

The pH along the root was automatically measured from 10 to 25 pixels off the root surface using custom Python scripts (ATR v5, https://sourceforge.net/projects/atr-along-the-root/, See Fig.S1f) averaging bins of 15*20 pixels from the fluorescence intensity of both 405 and 488nM channels.

Root angles over time were automatically measured using ACORBA v1.2 (Serre and Fendrych 2022).

### Statistical analysis and graphics

Statistical analyses were performed using the R software (R v4.0.2 and RStudio v1.3.1073). Boxplots represent the median and the first and third quartiles, and the whiskers extend to data points <1.5 interquartile range away from the first or third quartile; all data points are shown as individual dots. The results of the statistical tests are compiled in Supplemental statistics.

Graphics were created with the R software or in Python (v3.8) with the Seaborn plugin (v0.11.2, https://doi.org/10.21105/joss.03021) and aesthetic modifications of the graphs (fonts, size) were modified in Inkscape (v1.0).

## Supporting information

Supplemental Statistics

## Author contribution

DW assessed the already available pH-sensitive dyes and developed the method to use FS as a pH sensor. NCBS used the FS method to screen the pH profile of various genotypes in the study and developed the inducible lines. EM performed the line crossings and lab support. AJ performed the immunolabeling experiments. PV developed the CNGC14 complemented lines. SMD performed high temporal resolution imaging of root surface pH. DW, NCBS, JP, GG and MF wrote the manuscript.

## Data and source code availability

All raw and analyzed data and examples of scripts for measurements, statistics with R or Python will be available at Zenodo. Custom programs used in this article are available on their respective online repositories: ACORBA (https://sourceforge.net/projects/acorba/), ATR (https://sourceforge.net/projects/atr-along-the-root/).

## Acknowledgments

We would like to thank Mark Estelle for sharing the seeds; Karel Müller and Tomáš Moravec for help with molecular cloning; Mayuri Sadoine and Wolf Fommer for access to their spectrofluorometer and technical guidance.

## Supplemental Figures

**Figure S1:**
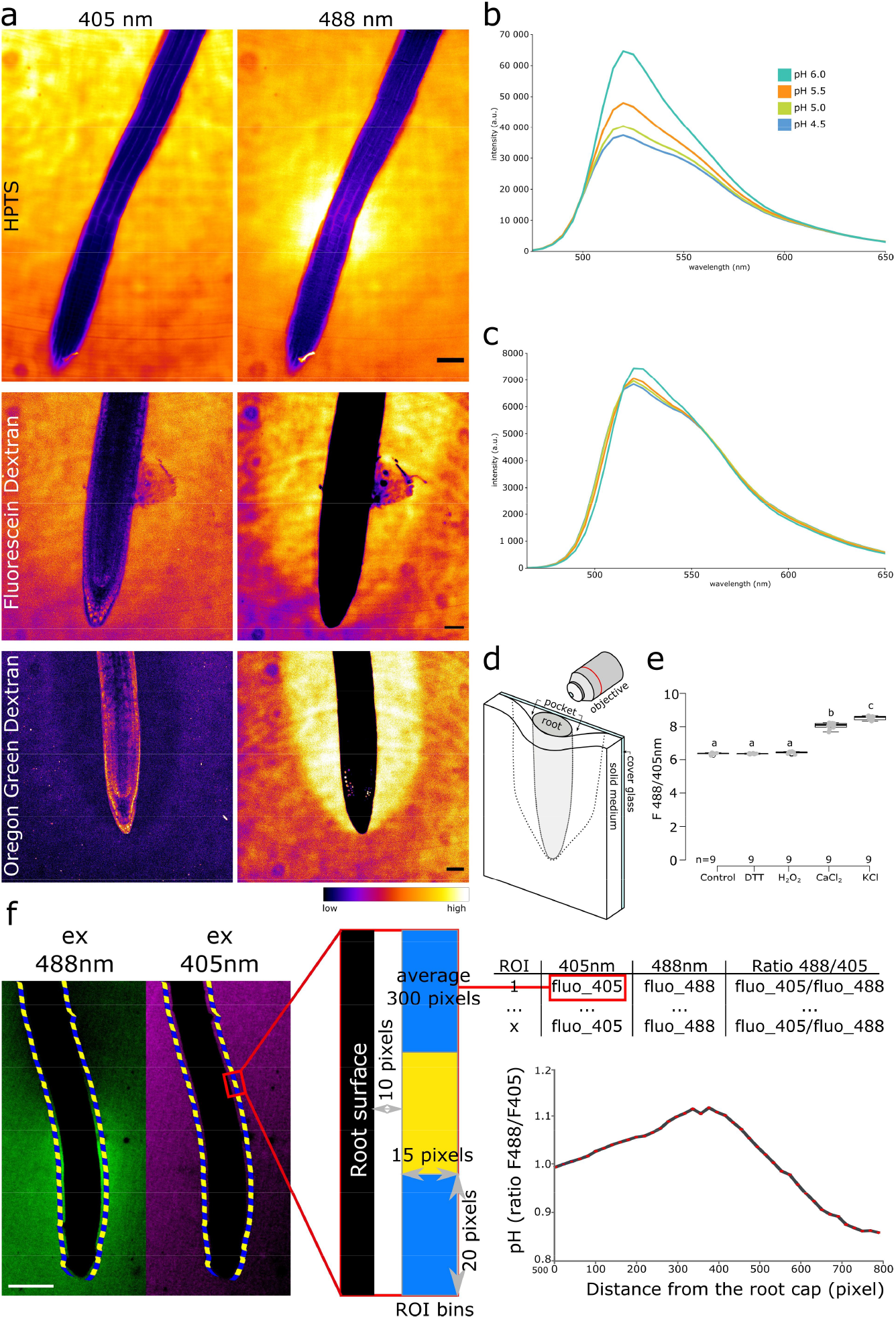
(a) Root surface pH imaged by HPTS, Fluorescein Dextran and Oregon Green Dextran. For all dyes, the reference excitation channel (405 nm) and the pH-responsive excitation channel (488 nm) are shown. HPTS shows an alkaline domain at the transition zone, while Fluorescein Dextran, and Oregon Green Dextran accumulate in a “pocket” (see (d)), preventing the observation of longitudinal pH profile. Emission was recorded at 500-550 nm. Col-0 line was used for HPTS and Oregon Green Dextran, DII-Venus line was used for Oregon Green Dextran. Scale bars = 50 μm. (b, c) pH-dependent FS spectra in liquid ½ MS. (b) Excitation by ν_EX_= 458 nm. (c) Excitation by ν_EX_= 405 nm. (d) Schematic representation of a root mounted into a microscopy chamber on a solid growth medium. Not drawn to scale. (e) FS F_488/405_ excitation ratio response to redox status and cations in MES buffered agar. Concentrations: 50 mM H_2_O_2_, 1 mM DTT, 10 mM CaCl_2_, 100 mM KCl. (f) The principle of the <u>A</u>long <u>T</u>he <u>R</u>oot (ATR) surface pH measurements by the FS F_488/405_ excitation ratio. The root surface is automatically recognized, creating a defined region of interest (ROI) in a 10-pixel distance from the surface. Fluorescence intensities in 488 and 405 nm excitation channels are measured in the ROIs. The data is binned and the FS F_488/405_ excitation ratio is calculated as a function of distance from the root tip. Scale bar = 100 μm.

**Figure S2:**
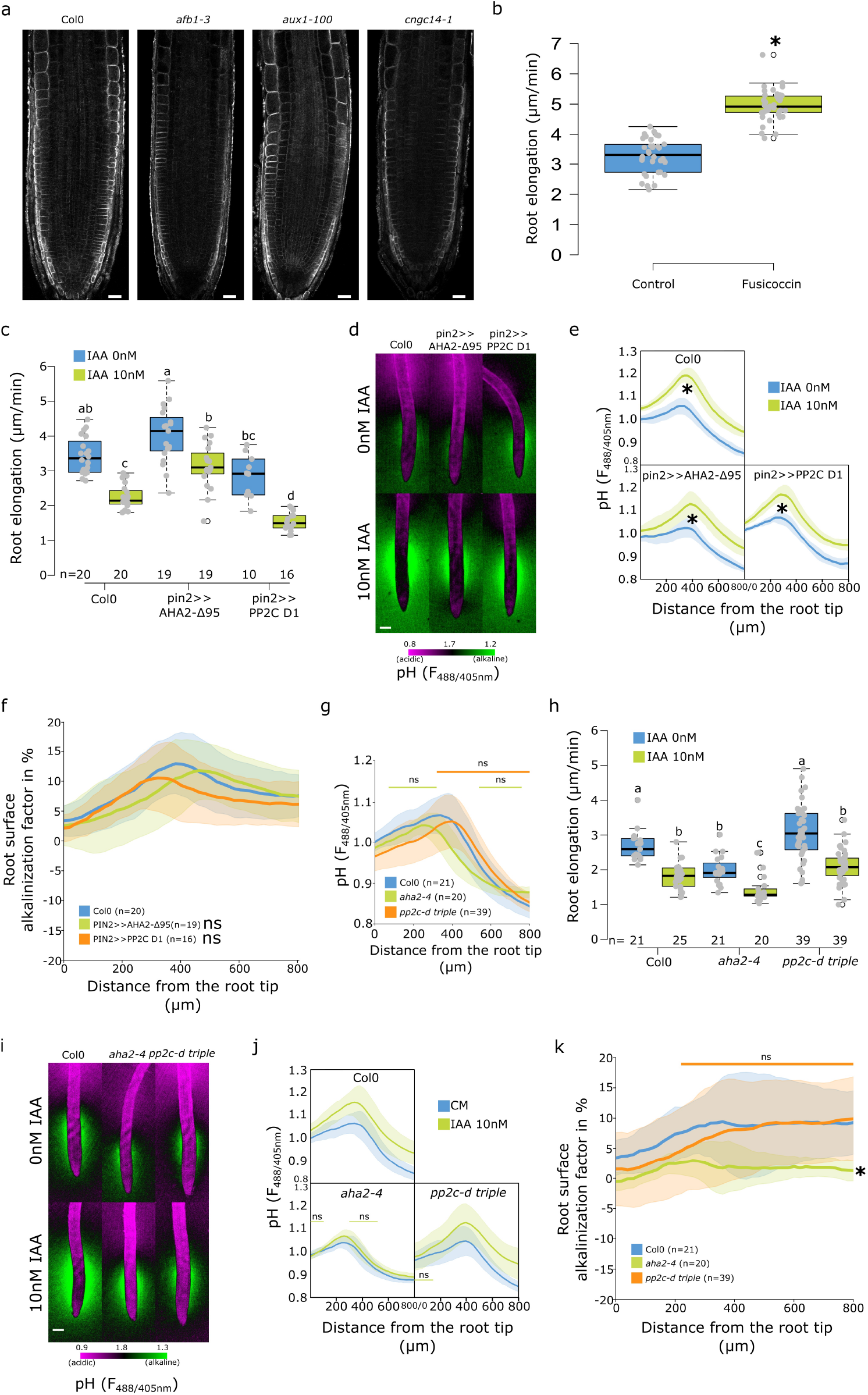
(a) Immunostaining of AHA proton pumps in Col-0, *afb1-3, aux1* and *cngc14-1* primary root tips. Scale bar = 20 μm. (b) Root elongation (μm/minute) of Col-0 treated with 0 or 2μM fusicoccin. Growth was measured over 40 min of imaging. (c-f) Roots of Col-0 and PIN2>>AHA2-Δ95 and PIN2>>PP2C-D1 lines after 4 hours induction by 5 μM estradiol and 20 minutes treatment with 0 or 10 nM IAA. (c) Root elongation (μm/minute) measured over 40 min. (d) Representative images of pH visualization by FS; scale bar = 100 μm. (e) Root surface pH quantification by FS F_488/405_ excitation ratio. (f) Comparison of root surface alkalinization induced by IAA expressed as a factor in % (ratio of individual IAA treated surface pH profile/average control profile × 100). (g-k) Roots of Col-0, *aha2-4* and *pp2c-d triple* mutants treated for 20 minutes with 0 or 10nM IAA. (g) Root surface FS F_488/405_ excitation ratio in control condition. (h) Root elongation (μm/minute) measured over 40 min. (i) Representative images of pH visualization by FS; scale bar = 100 μm). (j) Root surface FS F_488/405_ excitation ratio in response to IAA. (k) Root surface IAA alkalinization factor (as in (f). For b,c,h the letters on top of boxes correspond to the statistical groups. For e,f,g,j,k, ns: non significant statistical difference and *: p-value<.05.

**Figure S3:**
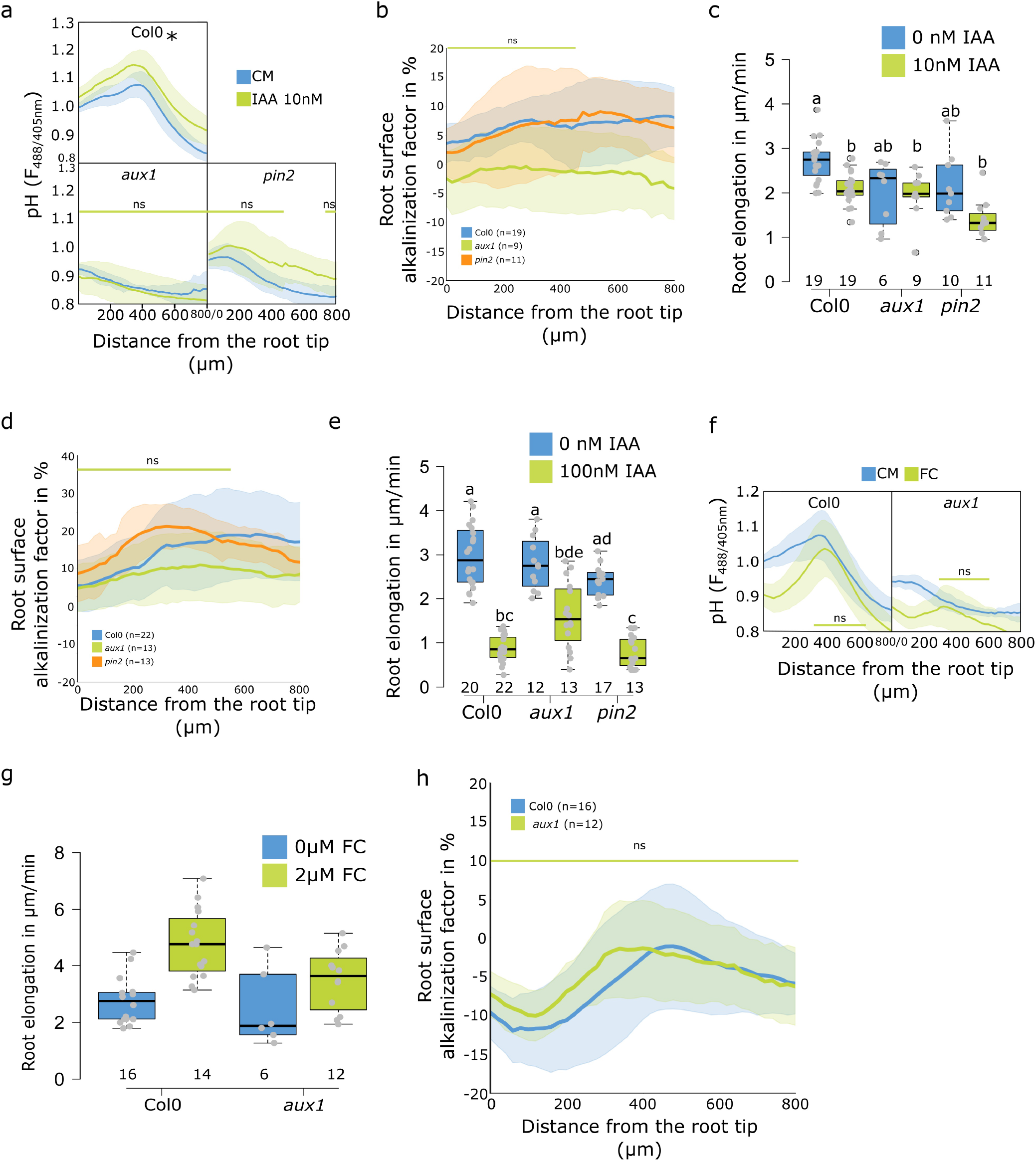
(a-c) Roots of Col-0, *aux1* and *pin2* mutants after 20 minutes treatment with 0 or 10nM IAA. (a) Root surface FS F_488/405_ excitation ratio. (b) Root surface alkalinization induced by IAA expressed as a factor in % (ratio of individual IAA treated surface pH profile/average control profile × 100). (c) Root elongation (μm/minute) measured over 40 min. (d,e) Roots of Col-0, *aux1* and *pin2* mutants after 20 minutes treatment with 0 or 100nM IAA. (d) Root surface alkalinization factor (as in (b)). (e) Root elongation (μm/minute) measured over 40 min. (f-h) Roots of Col-0 and *aux1* mutant after 20-min treatment with 0 or 2 μM fusicoccin. (f) Root surface FS F_488/405_ excitation ratio. (g) Root elongation (μm/minute) measured over 40 min. (h) Root surface alkalinization factor induced by 2 μM fusicoccin. For c,e,g, the letters on top of boxes correspond to the statistical groups. For a,b,d,f,h ns: non significant statistical difference and *: p-value<0.05.

**Figure S4:**
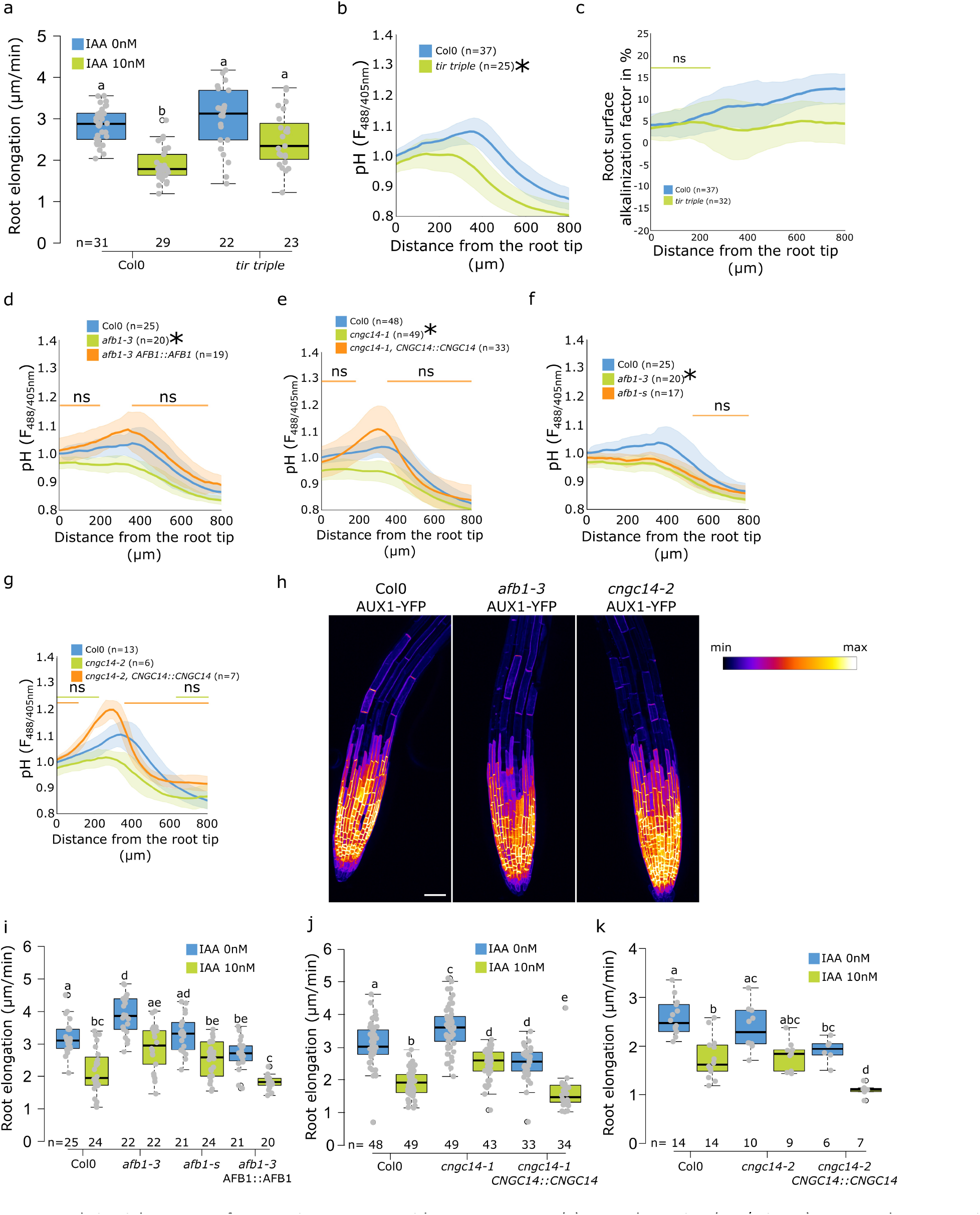
(a-c) Roots of Col-0 and *tir triple* mutant after 20 min treatment with 0 or 10nM IAA. (a) Root elongation (μm/minute) measured over 40 min. (b) Root surface FS F_488/405_ excitation ratio profiles in control condition. (c) Root surface alkalinization factor induced by IAA (ratio of individual IAA treated surface pH profile/average control profile × 100). (d) Comparison of root surface FS F_488/405_ excitation ratio profiles of Col-0, *afb1-3* mutant and AFB1::AFB1-mCitrine/*afb1-3* complemented line; comparison of genotypes from Fig.4d. (e) Comparison of root surface FS F_488/405_ excitation ratio profiles of Col-0, *cngc14-1* mutant and complemented line CNGC14::CNGC14-mVenus/*cngc14-1*; comparison of genotypes from Fig.4f. (f) Comparison of root surface FS F_488/405_ excitation ratio profiles of Col-0, *afb1-1s* and *afb1-3* mutants in control conditions. (g) Root surface FS F_488/405_ excitation ratio profiles of Col-0, *cngc14-2* mutant and the CNGC14::CNGC14-mVenus/*cngc14-2* complemented line. (h) localization of AUX1-YFP in the indicated mutant lines, scale bar = 50 μm. (i-k) Root elongation rates (μm/minute) in control and IAA-treated roots measured over 40 min in the indicated lines. For a,i,j,k the letters on top of boxes correspond to the statistical groups. For b,c,d,e,f,g ns= non significant statistical difference and *:p-value<0.05.

**Figure S5:**
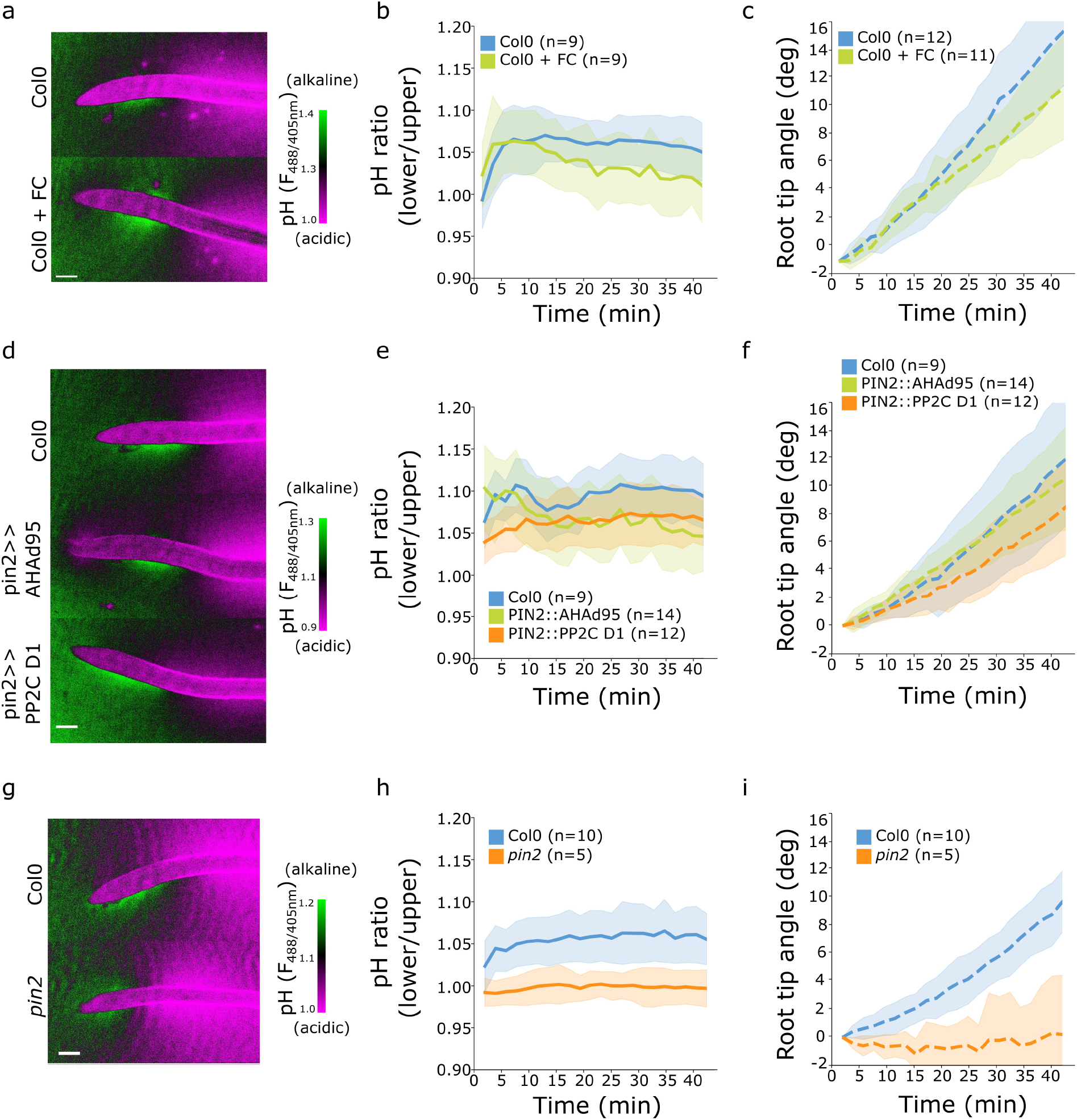
Surface pH dynamics and root tip bending angle during gravitropic responses of indicated lines and treatments. (a,d,g) Representative images of pH visualization by FS taken 40 minutes after gravistimulation. (b,e,h) Quantification of the FS F_488/405_ excitation ratio of lower/upper root transition zones. (c,f,i) Root tip angle dynamics as a function of time after gravistimulation. (a-c) Col-0 treated for 20 minutes with 0 or 2μM fusicoccin. (d-f) Col-0, PIN2>>AHA2-Δ95 and PIN2>>PP2C-D1 lines induced for 4h with 5μM estradiol. (g-i) Col-0 and *pin2* mutants. Scale bars = 100 μm.

## References

Ballio A, Chain EB, De Leo P, Erlanger BF, Mauri M, Tonolo A. 1964. Fusicoccin: a New Wilting Toxin produced by Fusicoccum amygdali Del. Nature 203: 297–297.

Band LR, Wells DM, Fozard JA, Ghetiu T, French AP, Pound MP, Wilson MH, Yu L, Li W, Hijazi HI, et al. 2014. Systems analysis of auxin transport in the Arabidopsis root apex. Plant Cell 26: 862–875.

Barbez E, Dünser K, Gaidora A, Lendl T, Busch W. 2017. Auxin steers root cell expansion via apoplastic pH regulation in Arabidopsis thaliana. Proc Natl Acad Sci 201613499.

Beemster GT, Baskin TI. 1998. Analysis of cell division and elongation underlying the developmental acceleration of root growth in Arabidopsis thaliana. Plant Physiol 116: 1515–26.

Behera S, Zhaolong X, Luoni L, Bonza MC, Doccula FG, De Michelis MI, Morris RJ, Schwarzländer M, Costa A. 2018. Cellular Ca2+ Signals Generate Defined pH Signatures in Plants. Plant Cell 30: 2704–2719.

Behrens HM, Weisenseel MH, Sievers A. 1982. Rapid Changes in the Pattern of Electric Current around the Root Tip of Lepidium sativum L. following Gravistimulation. Plant Physiol 70: 1079–83.

Bindels DS, Haarbosch L, van Weeren L, Postma M, Wiese KE, Mastop M, Aumonier S, Gotthard G, Royant A, Hink MA, et al. 2017. mScarlet: a bright monomeric red fluorescent protein for cellular imaging. Nat Methods 14: 1–12.

Björkman T, Leopold AC. 1987. An electric current associated with gravity sensing in maize roots. Plant Physiol 84: 841–6.

Braidwood L, Breuer C, Sugimoto K. 2014. My body is a cage: mechanisms and modulation of plant cell growth. New Phytol 201: 388–402.

Brunoud G, Wells DM, Oliva M, Larrieu A, Mirabet V, Burrow AH, Beeckman T, Kepinski S, Traas J, Bennett MJ, et al. 2012. A novel sensor to map auxin response and distribution at high spatiotemporal resolution. Nature 482: 103–6.

Campilho A, Garcia B, Toorn HVD, Wijk H V, Campilho A, Scheres B. 2006. Time-lapse analysis of stem-cell divisions in the Arabidopsis thaliana root meristem. Plant J 48: 619–27.

Clough SJ, Bent AF. 1998. Floral dip: a simplified method for Agrobacterium-mediated transformation of Arabidopsis thaliana. Plant J 16: 735–43.

Collings DA, White RG, Overall RL. 1992. Ionic Current Changes Associated with the Gravity-Induced Bending Response in Roots of Zea mays L. Plant Physiol 100: 1417–26.

Dharmasiri N, Dharmasiri S, Weijers D, Lechner E, Yamada M, Hobbie L, Ehrismann JS, Jürgens G, Estelle M. 2005. Plant development is regulated by a family of auxin receptor F box proteins. Dev Cell 9: 109–19.

Dindas J, Scherzer S, Roelfsema MRG, von Meyer K, Müller HM, Al-Rasheid KAS, Palme K, Dietrich P, Becker D, Bennett MJ, et al. 2018. AUX1-mediated root hair auxin influx governs SCFTIR1/AFB-type Ca2+ signaling. Nat Commun 9: 1174.

Dolan L, Janmaat K, Willemsen V, Linstead P, Poethig S, Roberts K, Scheres B. 1993. Cellular organisation of the Arabidopsis thaliana root. Development 119: 71–84.

Dubey SM, Serre NBC, Oulehlová D, Vittal P, Fendrych M. 2021. No Time for Transcription-Rapid Auxin Responses in Plants. Cold Spring Harb Perspect Biol 13: a039891.

Dusek J, Plchova H, Cerovska N, Poborilova Z, Navratil O, Kratochvilova K, Gunter C, Jacobs R, Hitzeroth II, Rybicki EP, et al. 2020. Extended Set of GoldenBraid Compatible Vectors for Fast Assembly of Multigenic Constructs and Their Use to Create Geminiviral Expression Vectors. Front Plant Sci 11: 522059.

Evans ML, Ishikawa H, Estelle MA. 1994. Responses ofArabidopsis roots to auxin studied with high temporal resolution: Comparison of wild type and auxin-response mutants. Planta 194: 215–222.

Evans ML, Mulkey TJ, Vesper MJ. 1980. Auxin Action on Proton Influx in Corn Roots and its Correlation with Growth. Planta 148: 510–512.

Falhof J, Pedersen JT, Fuglsang AT, Palmgren M. 2015. Plasma membrane H+-ATPase regulation in the center of plant physiology. Mol Plant 9: 323–337.

Fendrych M, Akhmanova M, Merrin J, Glanc M, Hagihara S, Takahashi K, Uchida N, Torii KU, Friml J. 2018. Rapid and reversible root growth inhibition by TIR1 auxin signalling. Nat plants 4: 453–459.

Friml J, Gallei M, Gelová Z, Johnson A, Mazur E, Monzer A, Rodriguez L, Roosjen M, Verstraeten I, Živanović BD, et al. 2022. ABP1-TMK auxin perception for global phosphorylation and auxin canalization. Nature 24.

Geilfus C-M, Mühling KH. 2011. Real-Time Imaging of Leaf Apoplastic pH Dynamics in Response to NaCl Stress. Front Plant Sci 2: 13.

Grieneisen VA, Xu J, Marée AFM, Hogeweg P, Scheres B. 2007. Auxin transport is sufficient to generate a maximum and gradient guiding root growth. Nature 449: 1008–1013.

Großeholz R, Wanke F, Rohr L, Glöckner N, Rausch L, Scholl S, Scacchi E, Spazierer A-J, Shabala L, Shabala S, et al. 2022. Computational modeling and quantitative physiology reveal central parameters for brassinosteroid-regulated early cell physiological processes linked to elongation growth of the Arabidopsis root. Elife 11.

Hager A. 2003. Role of the plasma membrane H+-ATPase in auxin-induced elongation growth: historical and new aspects. J Plant Res 116: 483–505.

Haruta M, Burch HL, Nelson RB, Barrett-Wilt G, Kline KG, Mohsin SB, Young JC, Otegui MS, Sussman MR. 2010. Molecular characterization of mutant Arabidopsis plants with reduced plasma membrane proton pump activity. J Biol Chem 285: 17918–29.

Haruta M, Sussman MR. 2012. The effect of a genetically reduced plasma membrane protonmotive force on vegetative growth of Arabidopsis. Plant Physiol 158: 1158–71.

Haruta M, Tan LX, Bushey DB, Swanson SJ, Sussman MR. 2018. Environmental and Genetic Factors Regulating Localization of the Plant Plasma Membrane H+-ATPase. Plant Physiol 176: 364–377.

Iwabuchi A, Yano M, Shimizu H. 1989. Development of extracellular electric pattern aroundLepidium roots: its possible role in root growth and gravitropism. Protoplasma 148: 94–100.

Jacobsen AGR, Jervis G, Xu J, Topping JF, Lindsey K. 2021. Root growth responses to mechanical impedance are regulated by a network of ROS, ethylene and auxin signalling in Arabidopsis. New Phytol 231: 225–242.

Li L, Verstraeten I, Roosjen M, Takahashi K, Rodriguez L, Merrin J, Chen J, Shabala L, Smet W, Ren H, et al. 2021. Cell surface and intracellular auxin signalling for H+ fluxes in root growth. Nature 599: 273–277.

Lin W, Zhou X, Tang W, Takahashi K, Pan X, Dai J, Ren H, Zhu X, Pan S, Zheng H, et al. 2021. TMK-based cell-surface auxin signalling activates cell-wall acidification. Nature 599: 278–282.

Lintilhac PM. 2014. The problem of morphogenesis: unscripted biophysical control systems in plants. Protoplasma 251: 25–36.

Liu S, Strauss S, Adibi M, Mosca G, Yoshida S, Dello Ioio R, Runions A, Andersen TG, Grossmann G, Huijser P, et al. 2022. Cytokinin promotes growth cessation in the Arabidopsis root. Curr Biol 32: 1974–1985.e3.

Lomax TL, Mehlhorn RJ, Briggs WR. 1985. Active auxin uptake by zucchini membrane vesicles: Quantitation using ESR volume and DELTApH determination. Proc Natl Acad Sci U S A 82: 6541–6545.

Luschnig C, Gaxiola RA, Grisafi P, Fink GR. 1998. EIR1, a root-specific protein involved in auxin transport, is required for gravitropism in Arabidopsis thaliana. Genes Dev 12: 2175–87.

Lüthen H, Böttger M. 1988. Kinetics of proton secretion and growth in maize roots: action of various plant growth effectors. Plant Sci 54: 37–43.

Mancuso S, Marras AM, Magnus V, Baluška F. 2005. Noninvasive and continuous recordings of auxin fluxes in intact root apex with a carbon nanotube-modified and self-referencing microelectrode. Anal Biochem 341: 344–351.

Martín-Barranco A, Thomine S, Vert G, Zelazny E. 2021. A quick journey into the diversity of iron uptake strategies in photosynthetic organisms. Plant Signal Behav 16: 1975088.

Martinière A, Bassil E, Jublanc E, Alcon C, Reguera M, Sentenac H, Blumwald E, Paris N. 2013. In vivo intracellular pH measurements in tobacco and Arabidopsis reveal an unexpected pH gradient in the endomembrane system. Plant Cell 25: 4028–43.

Monshausen GB, Miller ND, Murphy AS, Gilroy S. 2011. Dynamics of auxin-dependent Ca2+ and pH signaling in root growth revealed by integrating high-resolution imaging with automated computer vision-based analysis. Plant J 65: 309–18.

Monshausen GB, Zieschang HE, Sievers A. 1996. Differential proton secretion in the apical elongation zone caused by gravistimulation is induced by a signal from the root cap. Plant Cell Environ 19: 1408–14.

Moreau H, Gaillard I, Paris N. 2022. Genetically encoded fluorescent sensors adapted to acidic pH highlight subdomains within the plant cell apoplast. J Exp Bot.

Mulkey TJ, Evans ML. 1981. Geotropism in corn roots: evidence for its mediation by differential Acid efflux. Science 212: 70–1.

Müller A, Guan C, Gälweiler L, Tänzler P, Huijser P, Marchant A, Parry G, Bennett M, Wisman E, Palme K. 1998. AtPIN2 defines a locus of Arabidopsis for root gravitropism control. EMBO J 17: 6903–11.

Pacheco-Villalobos D, Díaz-Moreno SM, van der Schuren A, Tamaki T, Kang YH, Gujas B, Novak O, Jaspert N, Li Z, Wolf S, et al. 2016. The Effects of High Steady State Auxin Levels on Root Cell Elongation in Brachypodium. Plant Cell 28: 1009–24.

Pacifici E, Di Mambro R, Dello Ioio R, Costantino P, Sabatini S. 2018. Acidic cell elongation drives cell differentiation in the Arabidopsis root. EMBO J 37: e99134.

Prigge MJ, Platre M, Kadakia N, Zhang Y, Greenham K, Szutu W, Pandey BK, Bhosale RA, Bennett MJ, Busch W, et al. 2020. Genetic analysis of the Arabidopsis TIR1/AFB auxin receptors reveals both overlapping and specialized functions. Elife 9: 1–28.

Qi L, Kwiatkowski M, Chen H, Hoermayer L, Sinclair S, Zou M, Del Genio CI, Kubeš MF, Napier R, Jaworski K, et al. 2022. Adenylate cyclase activity of TIR1/AFB auxin receptors in plants. Nature 611: 133–138.

Rayle DL, Cleland RE. 1992. The Acid Growth Theory of auxin-induced cell elongation is alive and well. Plant Physiol 99: 1271–4.

Ren H, Park MY, Spartz AK, Wong JH, Gray WM. 2018. A subset of plasma membrane-localized PP2C.D phosphatases negatively regulate SAUR-mediated cell expansion in Arabidopsis. PLoS Genet 14: e1007455.

Retzer K, Akhmanova M, Konstantinova N, Malínská K, Leitner J, Petrášek J, Luschnig C. 2019. Brassinosteroid signaling delimits root gravitropism via sorting of the Arabidopsis PIN2 auxin transporter. Nat Commun 10: 5516.

Rosario LM, Rojas E. 1986. Modulation of K+ conductance by intracellular pH in pancreatic beta-cells. FEBS Lett 200: 203–9.

Rubery PH, Sheldrake a R. 1973. Effect of pH and surface charge on cell uptake of auxin. Nat New Biol 244: 285–8.

Sarrion-Perdigones A, Vazquez-Vilar M, Palací J, Castelijns B, Forment J, Ziarsolo P, Blanca J, Granell A, Orzaez D. 2013. GoldenBraid 2.0: a comprehensive DNA assembly framework for plant synthetic biology. Plant Physiol 162: 1618–31.

Sauer M, Paciorek T, Benková E, Friml J. 2006. Immunocytochemical techniques for whole-mount in situ protein localization in plants. Nat Protoc 1: 98–103.

Seksek O, Biwersi J, Verkman AS. 1995. Direct measurement of trans-Golgi pH in living cells and regulation by second messengers. J Biol Chem 270: 4967–70.

Serre NBC, Fendrych M. 2022. ACORBA: Automated workflow to measure Arabidopsis thaliana root tip angle dynamics. Quant Plant Biol 3: e9.

Serre NBC, Kralík D, Yun P, Slouka Z, Shabala S, Fendrych M. 2021. AFB1 controls rapid auxin signalling through membrane depolarization in Arabidopsis thaliana root. Nat plants 7: 1229–1238.

Shih HW, Depew CL, Miller ND, Monshausen GB. 2015. The cyclic nucleotide-gated channel CNGC14 regulates root gravitropism in arabidopsis thaliana. Curr Biol 25: 3119–3125.

Shih HW, Miller N, Dai C, Spalding E, Monshausen G. 2014. The Receptor-like Kinase FERONIA Is Required for Mechanical Signal Transduction in Arabidopsis Seedlings. Curr Biol 24: 1887–1892.

Siao W, Coskun D, Baluška F, Kronzucker HJ, Xu W. 2020. Root-Apex Proton Fluxes at the Centre of Soil-Stress Acclimation. Trends Plant Sci 25: 794–804.

Spartz AK, Ren H, Park MY, Grandt KN, Lee SH, Murphy AS, Sussman MR, Overvoorde PJ, Gray WM. 2014. SAUR Inhibition of PP2C-D Phosphatases Activates Plasma Membrane H+-ATPases to Promote Cell Expansion in Arabidopsis. Plant Cell 26: 2129–2142.

Staal M, De Cnodder T, Simon D, Vandenbussche F, Van der Straeten D, Verbelen J, Elzenga T, Vissenberg K. 2011. Apoplastic alkalinization is instrumental for the inhibition of cell elongation in the Arabidopsis root by the ethylene precursor 1-aminocyclopropane-1-carboxylic acid. Plant Physiol 155: 2049–2055.

Stanković B. 2006. Electrophysiology of Plant Gravitropism. In Plant Electrophysiology, pp. 423–436, Springer Berlin Heidelberg, Berlin, Heidelberg.

Swarup R, Kargul J, Marchant A, Zadik D, Rahman A, Mills R, Yemm A, May S, Williams L, Millner P, et al. 2004. Structure-function analysis of the presumptive Arabidopsis auxin permease AUX1. Plant Cell 16: 3069–83.

Swarup R, Kramer EM, Perry P, Knox K, Leyser HMO, Haseloff J, Beemster GTSS, Bhalerao R, Bennett MJ. 2005. Root gravitropism requires lateral root cap and epidermal cells for transport and response to a mobile auxin signal. Nat Cell Biol 7: 1057–65.

Verbelen J-P, De Cnodder T, Le J, Vissenberg K, Baluska F. 2006. The Root Apex of Arabidopsis thaliana Consists of Four Distinct Zones of Growth Activities: Meristematic Zone, Transition Zone, Fast Elongation Zone and Growth Terminating Zone. Plant Signal Behav 1: 296–304.

Weisenseel MH, Becker HF, Ehlgötz JG. 1992. Growth, Gravitropism, and Endogenous Ion Currents of Cress Roots (Lepidium sativum L.) : Measurements Using a Novel Three-Dimensional Recording Probe. Plant Physiol 100: 16–25.

Weisenseel MH, Dorn A, Jaffe LF. 1979. Natural H Currents Traverse Growing Roots and Root Hairs of Barley (Hordeum vulgare L.). Plant Physiol 64: 512–8.

Weisenseel MH, Meyer AJ. 1997. Bioelectricity, gravity and plants. Planta 203: S98–106.

Wisniewska J, Xu J, Seifertová D, Brewer PB, Ruzicka K, Blilou I, Rouquié D, Benková E, Scheres B, Friml J. 2006. Polar PIN localization directs auxin flow in plants. Science 312: 883.

Yang Y, Hammes UZ, Taylor CG, Schachtman DP, Nielsen E. 2006. High-affinity auxin transport by the AUX1 influx carrier protein. Curr Biol 16: 1123–7.

Zieschang H, Kohler K, Sievers A. 1993. Changing proton concentrations at the surfaces of gravistimulated Phleum roots. Planta 190.

Zuo J, Niu QW, Chua NH. 2000. An estrogen receptor-based transactivator XVE mediates highly inducible gene expression in transgenic plants. Plant J 24: 265–273.

